# Spermatogonial stem cells and their differentiation roadmap throughout marmoset development

**DOI:** 10.64898/2026.04.02.713872

**Authors:** Juan Manuel Paturlanne, Sara Di Persio, Toni Lange, Amadeusz Odroniec, Jahnavi Bhaskaran, Daniela Fietz, Xiaolin Li, Marius Wöste, Joachim Wistuba, Alexander Siegfried Busch, Juan M. Vaquerizas, Gerd Meyer zu Hörste, Sarah Sandmann, Stefan Schlatt, Sandra Laurentino, Nina Neuhaus

## Abstract

Sperm transfer genetic information from one generation to the next. These cells originate from gonocytes, which are specified during early embryonal development. To date however, there is limited information on the transcriptional gatekeepers governing the key events leading to sperm production: transition of gonocytes out of pluripotency around the time of birth, formation of the spermatogonial compartment prior to puberty, and initiation of germ cell differentiation at the time of puberty. We address this knowledge gap by employing the marmoset monkey (*Callithrix jacchus*) as a model for human postnatal testicular development. We analysed the transcriptional profiles of ∼48,000 neonatal, pre-pubertal, pubertal, and adult testicular cells and correlated these with histomorphometric measurements. We uncovered a transcriptional state linking gonocytes and spermatogonia characterized by CITED2 expression. Moreover, we propose NANOS2 and DPPA4 as regulators of spermatogonial plasticity from pre-puberty onwards and identify molecular gatekeepers of male germ cell differentiation.

## Introduction

In primates, primordial germ cells (PGCs) are specified during early embryonic development via the activation of transcription factors relevant for pluripotency (OCT4, AP2γ) and the germ cell lineage (DAZL, NANOS3).^1^ Once PGCs enter the genital ridge and become enclosed by somatic Sertoli and peritubular cell precursors, they are referred to as gonocytes, which continue to express pluripotency and germ cell markers.^2^ Relevantly, in primates, including the human, as the gonocytes migrate towards the basement membrane of the seminiferous tubules, they lose the expression of pluripotency markers and are then referred to as spermatogonia.^3^

In recent years, a growing body of single cell RNA sequencing (scRNA-seq) data from human fetal to adult testicular tissues has progressively charted the transcriptional landscape of the human male germline.^3–5^ Studies consistently report that the adult human spermatogonial compartment is characterized by the co-existence of undifferentiated spermatogonial states showing expression of *PIWIL4* (State 0), *FGFR3* (State 0A), *NANOS2* (State 0B), and *NANOS3* (State 1).^3–5^ Among these states, PIWIL4^+^ spermatogonia are considered the most undifferentiated germ cells.^6^

Based on mouse data, PIWIL4 is indispensable for maintaining DNA integrity by silencing transposable elements via *de novo* DNA methylation in gonocytes,^7,8^ hence the formation of this spermatogonial substate is crucial. However, results from human fetal/infantile testes remain inconclusive. On the one hand, a direct transition of germ cells expressing pluripotency markers into State 0 spermatogonia has been suggested to occur without intermediate substates in fetal tissues.^9^ On the other hand, another study did not capture State 0 spermatogonia in neonatal testicular tissues,^3^ suggesting that the human germ cell differentiation trajectory from gonocytes to PIWIL4^+^ spermatogonia has not been fully captured yet. Prior to puberty, PIWIL4^+^ spermatogonia give rise to the different marker-based transcriptional States 0-1 in the growing spermatogonial compartment.^10^ However, the transcriptional pathways and gatekeepers ensuring the formation and expansion of this compartment remain largely unknown.

Knowledge of the regulatory networks guiding the germ cell differentiation trajectory, from gonocytes all the way to sperm, is still largely missing for primates. This knowledge gap has direct negative consequences, as it precludes the identification of novel causes for male infertility,^11^ the development of *in vitro* testis models,^12^ and research on testicular germ cell cancer, which is caused by persisting gonocytes and is rising in incidence worldwide.^13^

These knowledge gaps are best addressed using an animal model, allowing systematic analyses of distinct developmental stages. The marmoset monkey (*Callithrix jacchus*) is considered the most suitable animal model for early human male germ cell development, as its spermatogonial stem cell system resembles that of the human,^14,15^ which differs significantly from that of rodents.^16^ Moreover, at the time of birth, marmoset testicular tissues are relatively immature, corresponding to the late fetal period in the human and allowing capture of the gonocyte-to-spermatogonia transition.^17,18^ Hence, this non-human primate model opens the door to a developmental period virtually inaccessible for study in the human. However, to date, scRNA-seq data of different developmental timepoints from marmosets are still lacking.

In this study, we focused on the germ cell differentiation trajectory from the transition out of pluripotency, through the formation of the spermatogonial compartment, to the production of sperm. To delineate the cellular transitions as well as the transcriptional switches, we used testicular tissues from neonatal, pre-pubertal, pubertal, and adult marmoset monkeys and performed scRNA-seq, which we complemented by quantitative immunohistomorphometrical analyses in an extended cohort. We thereby uncovered the transcriptional changes driving male germ cell differentiation from gonocytes to sperm.

## Results

### Single-cell transcriptomes of marmoset testicular tissues throughout development

To capture the transcriptional changes governing male germ cell differentiation in primates, we generated single-cell transcriptomes from marmoset monkey testes. These analyses of neonatal, pre-pubertal, pubertal, and adult tissues were complemented by histomorphometric datasets (**Fig. 1A**). Evaluation of testicular tissue architecture revealed an increase in the area covered by seminiferous tubules from 21-35% in neonatal animals to 60-87% in adults (**Fig. 1B, S1A**; **Table S1**). The most prominent germ cell types were spermatogonia in neonatal and pre-pubertal, spermatocytes in pubertal, and elongated spermatids in adult testicular tissues (**Fig 1B; Fig. S1B**; **Table S1**).

**Figure 1:**
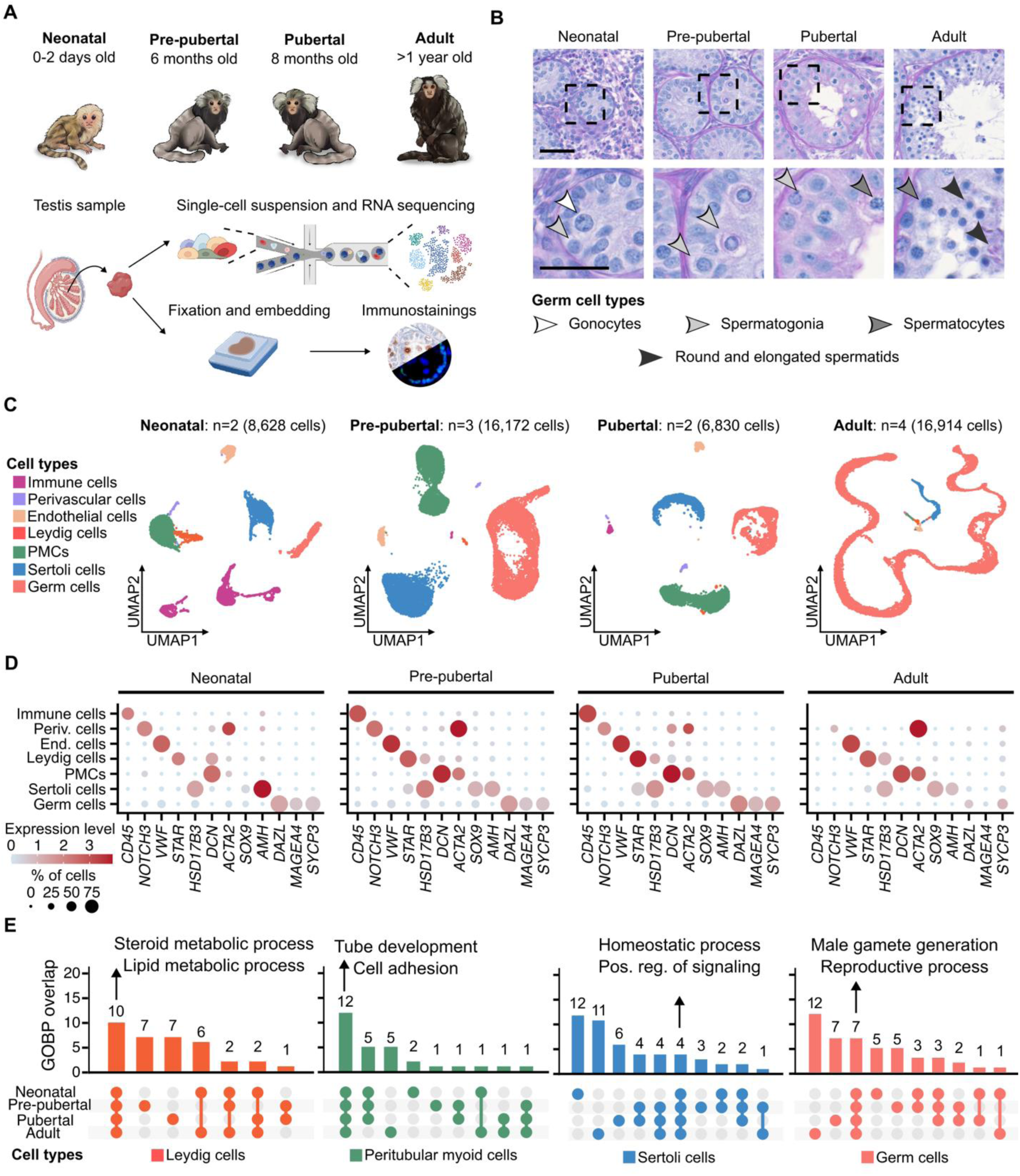
Cellular and transcriptional changes throughout marmoset postnatal testicular development. A) Experimental design. B) Representative histological micrographs of testicular tissues with arrowheads indicating the different male germ cell types. Scale bars = 50 μm. C) Uniform manifold approximation and projection (UMAP) plot of the single-cell RNA-sequencing (scRNA-seq) datasets. Differentially expressed genes (DEGs) are pro-vided in **Table S1**. D) Dotplots showing the expression level of selected marker genes used for testicular cell type assignments. E) UpSet plot depicting the overlap of the top 20 gene ontology terms related to biological processes (GOBP) between the developmental stages. Colors match the ones used to indicate cluster identity. GO terms and overlap results are provided in **Table S1**.

Following scRNA-seq and quality control, we captured transcriptomes of 48,194 cells, with a mean RNA content of 11,048 transcripts and 4,261 genes per cell (**Table S1**). Based on testicular cell type marker genes,^6^ we assigned the identities of clusters composed of immune (*CD45*), perivascular (*NOTCH3*), endothelial (*VWF*), Leydig (*STAR, HSD17B3*), peritubular myoid (PMCs; *DCN, ACTA2*), Sertoli (*SOX9, AMH*), and germ (*DAZL, MAGEA4, SYCP3*) cells (**Fig. 1C, 1D, S1C** and **S1D**).

To evaluate cell type-specific transcriptional profiles, we performed differential gene expression analysis at each developmental time point. In line with the assigned cell identities, functional annotation of differentially expressed genes (DEGs) revealed overlapping terms for the respective cell types at different developmental stages (**Fig. 1E**; **Table S1**). For instance, for Leydig cells these terms included ‘steroid metabolic processes’, and for germ cells ‘male gamete generation’.

We identified drastic changes in the proportions of immune and germ cells during development (**Fig. S1C**). Immune cells were most prominent in neonatal tissues (28%). Therefore, we re-clustered this cell population and, based on DEGs and published marker genes, assigned neutrophils, mast, T, and B cells, monocytes, and macrophages (**Fig. S1E** and **S1F**; **Table S1**).^19^ The presence of neutrophils and macrophages was further corroborated by co-immunofluorescence stainings demonstrating cells double-positive for CD45/PGLYRP1 (neutrophiles) and CD45/CD163 (macrophages) (**Fig. S1G** and **S1H**). The proportion of germ cells captured by scRNA-seq changed dynamically during development, constituting 6.9% of all cells in neonatal, 43.8% in pre-pubertal, 26.9% in pubertal, and finally 93.7% in adult tissues (**Fig. S1C; Table S1**). To further scrutinize the transcriptional changes along postnatal germ cell differentiation, we focused on the germ cells of each developmental stage individually.

### Formation of the spermatogonial compartment in neonatal testes

To unravel how the transcriptome changes during the gonocyte-to-spermatogonia transition and how this process is transcriptionally regulated, we subset and re-clustered 599 neonatal germ cells characterized by *DAZL* expression (**Fig. 2A** and **2B**; **Table S2**). Based on DEGs and selected marker genes, we assigned four out of the five clusters: MKI67^+^ gonocytes (*MKI67*, *OCT4*, *AP2y*), gonocytes (*OCT4*, *AP2y*), early spermatogonia (*TEX15*, *NKX6-2*), and PIWIL4^+^/State 0 spermatogonia (*PIWIL4, EGR4*) (**Table S2**). Unlike *OCT4* and *AP2y*, the marker gene *LIN28A* was expressed from gonocytes to early spermatogonia and *MAGEA4* in early and *PIWIL4*^+^ spermatogonia (**Fig. 2B**). The fifth cluster, which we refer to as the transitioning germ cells (**Fig. 2A**), consisted of cells bridging the gonocyte and spermatogonial subpopulations and differentially expressed *CITED2* and *DNAH5* (**Table S2**). The upregulation of the former is of particular interest, as CITED2 has been demonstrated to downregulate genes associated with pluripotency and self-renewal in embryonic stem cells.^20,21^

**Figure 2:**
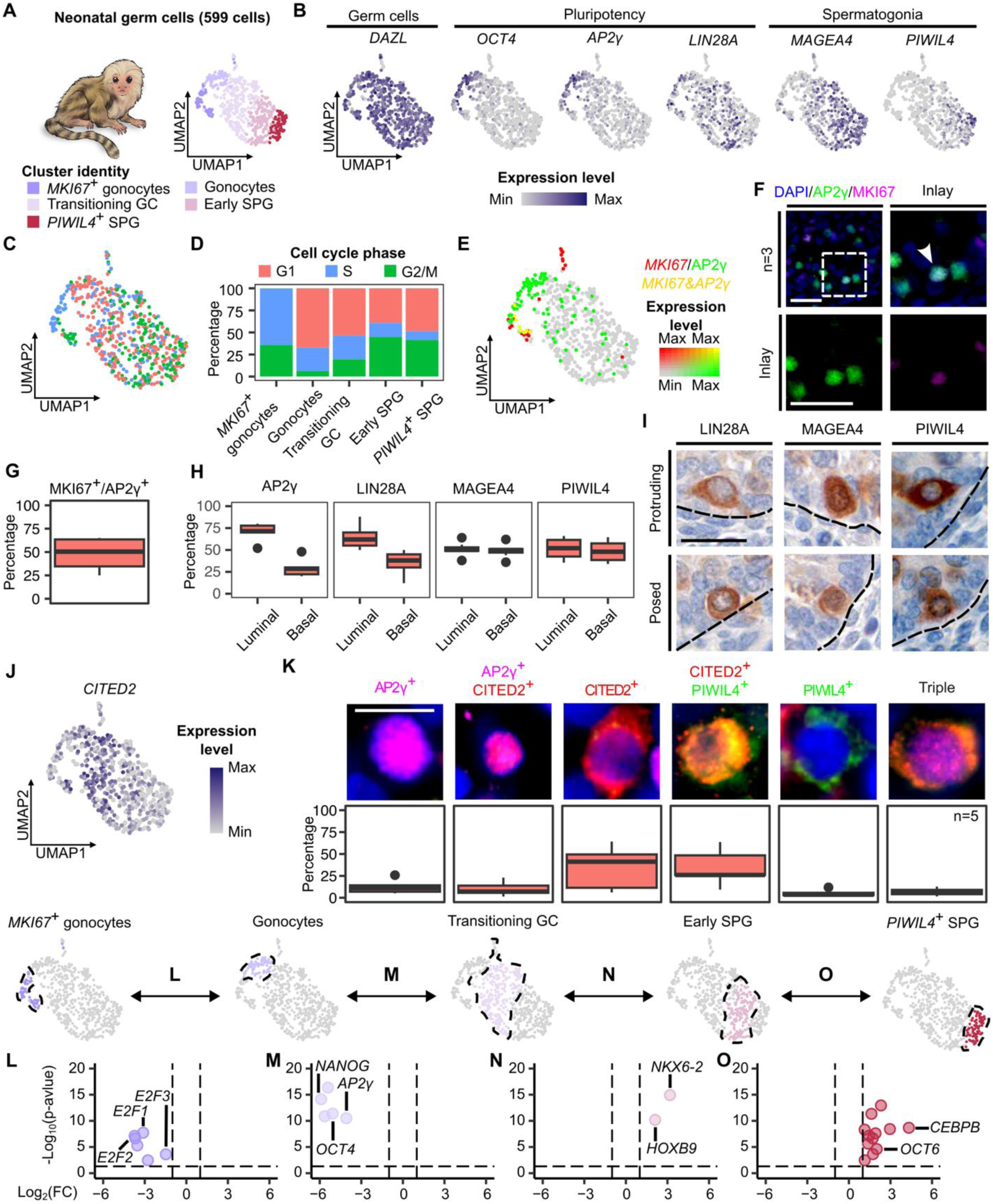
Gonocytes transition via upregulation of CITED2 into spermatogonia. A) UMAP plot of the neonatal germ cell subset (599 cells). DEGs are provided in **Table S2**. B) Feature plots showing expression of the marker genes used for cell cluster assignment. C) UMAP plot of neonatal germ cells categorized based on cell cycle analysis into G1, S, or G2/M phase. D) Barplots depicting the proportion of cells in G1, S, or G2/M phase per germ cell cluster. Results are provided in **Table S2**. E) Double feature plot showing the expression of the proliferation marker *MKI67* (red), gonocyte marker *AP2γ* (green), or both (yellow). F) Representative immunofluorescence micrographs of neonatal marmoset testicular tissues stained for AP2γ (green) and MKI67 (magenta) counterstained with DAPI (blue). The dashed square delineates the inlay area. White arrowhead points at double positive cells. Scale bar = 20 μm. G) Boxplot showing the percentage of double positive cells for MKI67 and AP2γ in neonatal tissues (n=3). Quantification results are provided in **Table S2**. H) Boxplots showing localization of marker-positive cells within the seminiferous tubules of neonatal marmoset tissues (n=7). Quantification results are provided in **Table S2**. I) Representative micrographs of neonatal marmoset testicular tissues stained for LIN28A, MAGEA4, or PIWIL4, showing cytoplasmic protrusions towards the tubular wall (top row) or located at the tubular wall (bottom row). Black dashed lines delineate the tubular wall. Scale bar = 20 μm. J) Feature plot of *CITED2*, a gene differentially expressed by Transitioning germ cells. K) Co-localization analysis of AP2γ (magenta), CITED2 (red), and PIWIL4 (green). Representative images of single, double, and triple positive cells (upper panel) and quantification results in neonatal marmoset testicular tissues (n=5), represented as box-plots. Quantification results are provided in **Table S2** L-O) Results of the differential regulon activity analysis. UMAP plots (top row) highlight the cluster comparisons and volcano plots depict respective results. Vertical dashed lines represent the fold change threshold (|log_2_(fold change)| = 1), and the horizontal dashed line represents the −log_10_(p-value) threshold (p-value = 0.05). Differentially active regulons are provided in **Table S2**.

Functional annotation of DEGs between the different germ cell types yielded terms including “cell cycle” for *MKI67*^+^ gonocytes (**Fig. S2A**; **Table S2**), supporting the co-existence of proliferating (*MKI67^+^*) and non-proliferating gonocytes (**Fig. 2A**). This was further supported by cell cycle analysis, which revealed that *MKI67*^+^ gonocytes consisted solely of cells in the S and G2/M phases (64.4% and 35.6%) (**Fig. 2C** and **2D**; **Table S2**). Moreover, we showed co-expression of *AP2γ* and *MKI67* at RNA level (**Fig. 2E**) and at protein level in 53.3% of AP2γ^+^ cells (**Fig. 2F** and **2G**; **Table S2**). For *PIWIL4*^+^ spermatogonia, functional annotation showed enrichment in “cell projection morphogenesis”, which includes genes involved in cell migration (**Fig. S2A**). To follow up on this, we quantified the positive cells with a luminal or basal localization within the seminiferous tubules and found 29.9% AP2γ^+^ cells located at the basal membrane compared to 49.1% of MAGEA4^+^ and 47.3% of PIWIL4^+^ cells (**Fig. 2H**). Additionally, in line with the functional enrichment results, LIN28A^+^, MAGEA4^+^, and PIWIL4^+^ cells showed cytoplasmic protrusions (**Fig. 2I**) as well as expression of relevant genes, such as *GFRA3* and *BAIAP2* (**Table S2**), which have been linked to cell migration.^22,23^

In line with the scRNA-seq data, quantitative histometrical evaluation of the germ cell compartment revealed lower numbers of cells positive for gonocyte markers per round tubule (0.4 OCT4^+^; 0.6 AP2γ^+^) compared to spermatogonial markers (1.7 MAGEA4^+^; 2.1 PIWIL4^+^) (**Fig. S2B**; **Table S2**). Moreover, the three samples with the fewest gonocytes contained higher numbers of PIWIL4^+^ cells (**Fig. S2B**; **Table S2**), reflecting differences in the timing of the transition from gonocytes to spermatogonia. Validating the existence of *CITED2*-expressing germ cells linking gonocytes to spermatogonia at protein level (**Fig. 2J**), we demonstrated subpopulations positive only for AP2γ (12.6%), CITED2 (34.4%), or PIWIL4 (6.2%), joined by subpopulations co-expressing these markers, reflecting the differentiation trajectory (**Fig. 2K**; **Table S2**). The gonocyte-to-spermatogonia transition is further reflected by the presence of cells expressing either LIN28A (43.6%) or MAGEA4 (6.1%) as well as a population immunopositive for both (50.2%) (**Fig. S2D**; **Table S2**).

To uncover the gene regulatory networks governing the transition from gonocytes to spermatogonia, we used SCENIC^24^ to identify regulons, *i.e.* groups of genes that are co-regulated by a transcription factor. Following identification of 35 overall differentially active regulons in the neonatal germ cell subset (**Fig. S2E**; **Table S2**), we performed sequential differential analysis along the gonocyte-to-spermatogonia transition to capture the transcriptional changes from one stage to the next (**Fig. 2L-O**; **Table S2**). While both gonocyte clusters shared activity of regulons associated with pluripotency (*NANOG, AP2γ, OCT4*), members of the family of E2F transcription factors (*E2F1*, *E2F2*, *E2F3*) were specifically active in the *MKI67*^+^ gonocytes (**Fig. 2L** and **S2E**; **Table S2**). These transcription factors play a major role in the G1/S phase transition,^25^ which aligns with the lack of cells in this cluster assigned to G1 stage in *MKI67*^+^ gonocytes (**Fig. 2D**; **Table S2**). Intriguingly, the cluster of transitioning germ cells was characterized by downregulation of the pluripotency cassette compared to gonocytes but did not display specific activity of regulons (**Fig. 2M**; **Table S2**). Following this regulatory ground state, we detected enhanced activity of *HOXB9* (10 regulated genes, including *ID4*) and *NKX6-2* (25 regulated genes, including *ID4* and *OCT6*) in early spermatogonia (**Fig. 2N**; **Table S2**). While *ID4* plays a role in the transition between spermatogonial stem cells and progenitor states in mouse,^26^ *OCT6* has a role in the regulation of GDNF-induced survival and self-renewal of mouse spermatogonial stem cells.^27^ The most notable regulatory change was detected in *PIWIL4*^+^ spermatogonia compared to early spermatogonia, with 12 activated regulons (*OCT6*, *CEBPB)* (**Fig. 2O**; **Table S2**).

In sum, we captured hitherto unknown cell states along the unidirectional germ cell differentiation trajectory, linking gonocytes to PIWIL4^+^ spermatogonia.

### The undifferentiated spermatogonial compartment in the pre-pubertal testis

To scrutinize the composition and transcriptional makeup of the undifferentiated spermatogonial compartment prior to puberty onset, we subset and re-clustered the 7,078 germ cells obtained from pre-pubertal testes (**Fig. 3A**; **Table S3**). These comprised 4 clusters of undifferentiated spermatogonia (*MAGEA4^+^, UTF1^+^*) (**Fig. 3A** and **3B**), whose identity we assigned based on DEGs. The two main undifferentiated spermatogonial substates were characterized by expression of *PIWIL4* (3,375 cells) and *NANOS3* (2,761 cells), respectively. When projected in the uniform manifold approximation and projection (UMAP), these substates formed a “roundabout” connected by an interface formed by a small cluster of cells differentially expressing *NANOS2* (490 cells), reflecting transcriptional plasticity (**Fig. 3A** and **3B**; **Table S3**). The remaining cluster, branching off the *NANOS3*^+^ spermatogonia, expressed *MKI67* and functional annotation of its DEGs substantiated that it is formed by proliferating cells (452 cells) (**Fig. S3A**; **Table S3**). This was further corroborated by cell cycle analysis, demonstrating that all *MKI67*^+^ spermatogonia were in S and G2/M-phase (50.2% and 49.8%, respectively) (**Fig. S3B**; **Table S3**) and co-expression of MKI67 in around 14% of NANOS3^+^ spermatogonia at protein level (**Fig. S3C** and **S3D**; **Table S3**). Among the DEGs of *NANOS3*^+^ spermatogonia were *CITED2* and *DPPA4* (**Table S3**). At protein level, we detected stable numbers of DPPA4-immunopositive spermatogonia per tubular cross-section throughout development (neonatal: 2; pre-pubertal: 1; pubertal: 1.4; adult: 1.8 DPPA4^+^) (**Fig. 3C**). Interestingly, *DPPA4* was mainly expressed at the interface between the *NANOS3*^+^ and *PIWIL4^+^* spermatogonial clusters (**Fig. 3D**). In line with this, double positive cells for DPPA4 and either NANOS3 or PIWIL4, and triple positive cells for all markers were detected at protein level (**Fig. 3E**), suggesting that this gene is associated with the transcriptional plasticity between the main spermatogonial substates.

**Figure 3:**
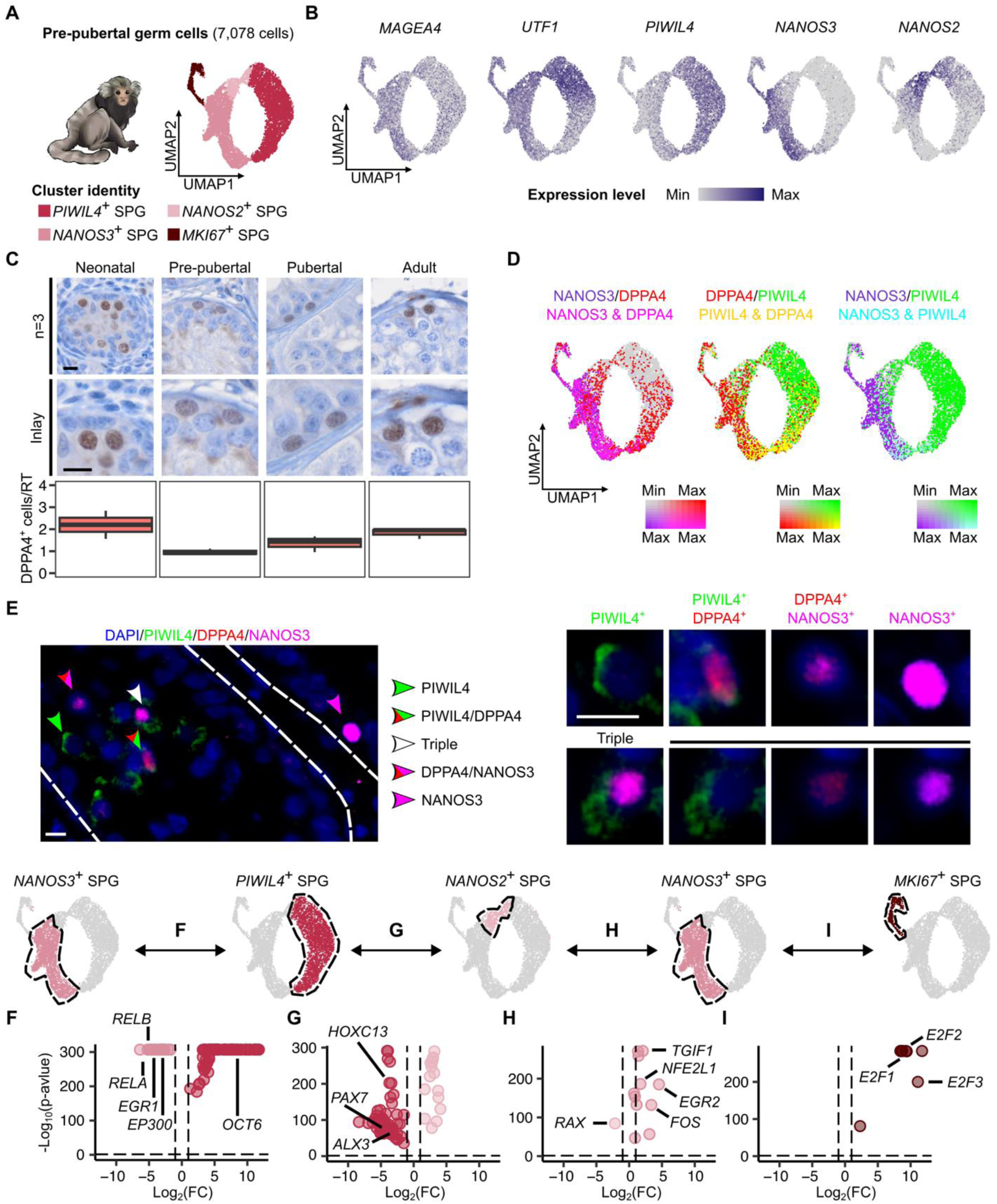
Transition of undifferentiated spermatogonia via *NANOS2* or *DPPA4* upregulation. A) UMAP plot of pre-pubertal germ cell subset (7,078 cells). DEGs are provided in **Table S3**. B) Feature plots showing the marker genes used for cell assignment. C) Representative histological micrographs of DPPA4 stainings throughout development (top row) and respective inlays (middle panel). Boxplots (bottom row) depict the quantification results of immunopositive cells per round (n=3 each). Scale bars = 20 μm and 10 μm in the inlays. D) Double feature plots highlighting co-expression of *NANOS3, DPPA4* and *PIWIL4* in the pre-pubertal germ cell subset. E) Representative immunofluorescence micrographs of pre-pubertal testicular tissue stained for NANOS3 (magenta), DPPA4 (red), and PIWIL4 (green). Arrow heads are color coded to indicate single, double, and triple positive cells. Representative triple positive cell in tissue section for PIWIL4, DPPA4, and NANOS3. Scale bar = 20 μm. F-I) Results of differential regulon activity analyses. UMAP plots (top row) highlight the clusters considered for each comparison and Vulcano plots show respective results. Vertical dashed lines represent the threshold at ±1 of log_2_(FoldChange), and the horizontal dashed line represents the threshold of −log_10_(p-value) for a p-value of 0.05. Differentially active regulons are provided in **Table S3**.

To uncover the transcriptional gatekeepers facilitating the transition between the spermatogonial states, we performed sequential differential regulon activity analysis (**Fig. 3F-I** and **S3E**; **Table S3**). 82 regulons were differentially active in the comparison between *NANOS3*^+^ (*EGR1, RELA, RELB,* and *EP300*) and *PIWIL4*^+^ spermatogonia (*OCT6*) (**Fig. 3F**; **Table S3**). EGR1 expression has been associated with cell proliferation and differentiation.^28^ Co-localization of EGR1 and MAGEA4 at single-cell RNA and protein levels validated the expression of EGR1 in marmoset spermatogonia throughout development (**Fig. S3F** and **S3G**; **Table S3**). We inferred the differential activation of 64 regulons in *PIWIL4*^+^ compared to *NANOS2*^+^ spermatogonia (**Fig. 3G**; **Table S3**). Of these, only three, *PAX7*, *ALX3,* and *HOXC13,* were exclusive to this comparison and not detected between *NANOS3*^+^ and *PIWIL4*^+^ spermatogonia (**Table S3**). Interestingly, *PAX7* has been reported to be expressed by a small subpopulation of undifferentiated spermatogonia surviving gonadotoxic therapy (chemo- and radiotherapy) in murine testes.^29^ From the 15 enriched regulons in *NANOS2*^+^ compared to *PIWIL4*^+^ spermatogonia, none were exclusive (**Table S3**). The comparison of *NANOS3*^+^ and *NANOS2*^+^ spermatogonia showed only one unique differentially active regulon in the latter, namely *RAX* (**Fig. 3H**; **Table S3**), which interestingly is predicted to upregulate *NANOS2* (**Table S3**). Four regulons were differentially active in *NANOS3*^+^ spermatogonia in the general comparison (**Fig. S3E**) and when compared to both *PIWIL4*^+^ and *NANOS2*^+^ spermatogonia (*TGIF1*, *EGR2*, *FOS,* and *NFE2L1)* (**Fig. 3F** and **3H**; **Table S3**), showing high specificity to this cluster. In contrast, apart from proliferation-related *E2F1, E2F2,* and *E2F3*, no specific regulons were upregulated in *NANOS3*^+^ versus *MKI67*^+^ spermatogonia (**Fig. 3I**; **Table S3**), corroborating their *NANOS3*^+^ spermatogonial identity.

In sum, undifferentiated spermatogonia are comprised of *PIWIL4* and *NANOS3*^+^ subpopulations that transition via upregulation of *NANOS2* and *DPPA4*. While *PIWIL4*^+^ spermatogonia express regulons such as *PAX7*, associated with the true stem cell identity, *NANOS3*^+^ cells include a proliferative cluster.

### Germ cell differentiation unleashed in the pubertal testis

To uncover the transcriptional changes accompanying the initiation of germ cell differentiation and meiotic entrance, we analyzed the pubertal germ cell subset consisting of 1,836 cells (**Fig. 4A**; **Table S4**). Based on DGE analysis, we identified undifferentiated spermatogonia (*UTF1, PIWIL4, NANOS2, NANOS3*), two clusters of differentiating spermatogonia (*KIT, STRA8, REC8*), pre- and leptotene spermatocytes (*MEIOB, TEX101*) (**Fig. 4A** and **4B**, **Table S4**). *KIT*^+^ spermatogonia showed upregulation of *RHOXF2B* (**Table S4**). Interestingly, in pre-pubertal germ cells, we also identified a small population of *NANOS3*^+^ spermatogonia expressing *RHOXF2B* (**Fig. 4C**), likely representing the starting point of differentiation. Indeed, we also identified immunopositive cells for RHOXF2B in seminiferous tubules of pre-pubertal and pubertal testis (**Fig. 4D**).

**Figure 4:**
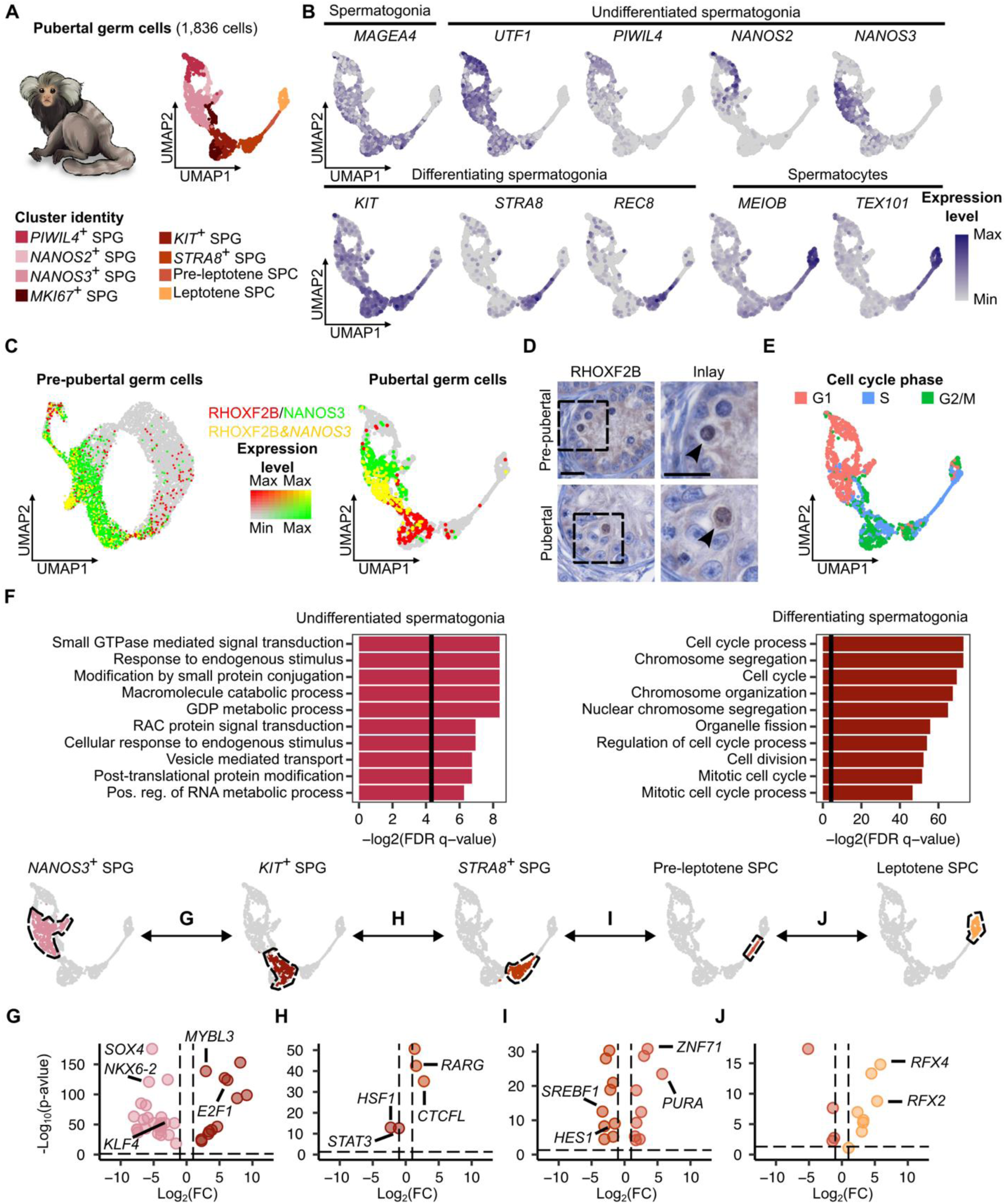
Germ cell differentiation is initiated by upregulation of RHOXF2B. A) UMAP plot of the pubertal germ cell subset (1,836 cells). DEGs are provided in **Table S4**. B) Feature plots depicting the marker genes used for cell assignment. C) Double feature plots showing expression of *NANOS3* (green) and *RHOXF2B* (red) in pre-pubertal (left) and pubertal (right) germ cell subsets. Double positive cells are colored in yellow. D) Representative histological micrographs of *RHOXF2B*^+^ cells in pre-pubertal and pubertal testicular tissues (arrow heads). Scale bars = 20 μm. E) UMAP plot of pubertal germ cells categorized based on the cell cycle analysis into G1, S, or G2/M phases. Results are provided in **Table S4**. F) GO terms associated with the top DEGs (up to 500) enriched in direct comparisons between undifferentiated spermatogonia (*PIWIL4*^+^ SPG, *NANOS2*^+^ SPG and *NANOS3*^+^ SPG), and differentiating spermatogonia (*KIT*^+^ SPG and *STRA8*^+^ SPG), in the pubertal germ cell subset. Bars depict the −log_2_ of the false discovery rate of each term and the vertical line corresponds to an FDR q-value = 0.05. GO terms are provided in **Table S4**. G-J) Results of the differential regulon activity analysis. UMAP plots in (top row) highlight the clusters considered for each comparison and volcano plots (bottom row) depict respective results. Vertical dashed lines represent the fold change threshold (|log_2_(fold change)| = 1), and the horizontal dashed line represents the −log_10_(p-value) threshold (p-value = 0.05). Differentially active regulons are provided in **Table S4**.

To scrutinize the biological meaning of the transcriptional differences between undifferentiated (*PIWIL4*^+^, *NANOS2*^+^, and *NANOS3*^+^) and differentiating spermatogonia (*KIT*^+^ and *STRA8*^+^), we performed differential gene expression analysis followed by functional annotation (**Fig. 4F**; **Table S4**). The top ten gene ontology terms enriched for differentiating spermatogonia were associated with cell cycle, including “chromosome organization and segregation”, “organelle fission”, and “mitotic cell cycle” (**Fig. 4F**; **Table S4**). Functional annotation of the cluster-specific DEGs further revealed that differentiating spermatogonia undergo proliferation, while (pre-)leptotene spermatocytes expressed genes (*MEIOB*, *MAJIN*) associated with “meiosis I cell cycle process” (**Fig. S4A**; **Table S4**). Indeed, cell cycle analysis indicated higher proliferation in differentiating spermatogonia, with most of the cells assigned to S or G2/M phase, respectively (*KIT*^+^: 53% and 46.7%; *STRA8*^+^: 45.9% and 45%) (**Fig. 4E** and **S4B**; **Table S4**). This is even more evident when compared to undifferentiated spermatogonia, with cells largely in G1 phase (*PIWIL4*^+^: 86.9%, *NANOS2*^+^: 98.9%, *NANOS3*^+^: 84%), except for the *MKI67*^+^ cluster, assigned to either S or G2/M phase (27.2% and 72.8%) (**Fig. 4E** and **S4B**; **Table S4**). Co-expression analyses at RNA and protein level underscored the co-existence of two proliferative (MKI67^+^) spermatogonial subpopulations among the NANOS3^+^ undifferentiated spermatogonia (11.5%) and the differentiating *KIT*^+^ spermatogonia, respectively (**Fig. S4C-F**). These findings indicate the expansion of both the undifferentiated and differentiating spermatogonial compartments from puberty onwards.

To dissect the transcriptional changes in male germ cell differentiation, we scrutinized the 132 regulons that were differentially active in pubertal germ cells in general (**Table S4**) and, among them, those that were sequentially activated from *NANOS3*^+^ spermatogonia up to leptotene spermatocytes (**Fig. 4G-J**; **Table S4**). *NANOS3*^+^ spermatogonia displayed differential activity of the regulons *KLF4*, *SOX4,* and *NKX6-2* (**Fig. 4G**; **Table S4**). Importantly, *SOX4* and *NKX6-2* were differentially active in all undifferentiated spermatogonial substates (**Fig. S4G**; **Table S4**). The main upregulated regulons in *KIT*^+^ spermatogonia (*MYBL3, E2F1*) belonged to proliferation programs (**Fig. 4G**; **Table S4**). Moving towards differentiation, comparison of *KIT*^+^ and *STRA8*^+^ spermatogonia revealed increased activity of regulons *HSF1* and *STAT3* in the former and of *CTCFL* and *RARG* in the latter cell population (**Fig. 4H**). STAT3 signaling leads to enhanced activity of TET enzymes,^30^ key mediators of epigenetic reprogramming while CTCFL has been reported as a master transcription factor regulating human spermatogenesis, with a peak in expression in *STRA8*^+^ differentiated spermatogonia,^31^ aligning with our results. When comparing *STRA8*^+^ spermatogonia to pre-leptotene spermatocytes, regulon activity analysis revealed *HES1 and SREBF1* in the former and *PURA* and ZNF71 in the latter (**Fig. 4I**). Finally, *RFX2* and *RFX4* were among the differentially active regulons of the leptotene spermatocytes (**Fig. 4J**; **Table S4**) and have been associated with meiosis.^32^

Our analyses highlight *SOX4* and *NKX6-2* as potential gatekeepers of undifferentiated spermatogonial identity, with the latter predicted to promote spermatogonial identity also in neonatal testes (**Fig. 2N**). Apart from that, at the pubertal stage, we report proliferation among *NANOS3^+^* undifferentiated spermatogonia as well as differentiating *KIT^+^* spermatogonia, the latter expanding the number of cells committing to meiosis.

### Full male germ cell differentiation and its regulation in the adult testis

In contrast to earlier developmental stages, in the adult testis we captured the whole scope of male germ cell differentiation (15,982 cells) (**Fig. 5A**; **Table S5**). This allowed us to uncover the transcriptional changes accompanying meiosis and spermiogenesis. We performed cell type assignment using DEGs specifically reported for adult marmosets.^33^ This allowed us to distinguish spermatogonia (MAGEA4) into undifferentiated (*UTF1*) and differentiating (*KIT*), spermatocytes into leptotene (*MEIOB*), zygotene (*TEX14*), pachytene (*CCNA1*), and diplotene (*ANKRD55*), and spermatids into early round (*SPACA3*), late round (*EQTN*), and elongated (*TNP1*) (**Fig. 5A** and **5B**; **Table S5**).

**Figure 5:**
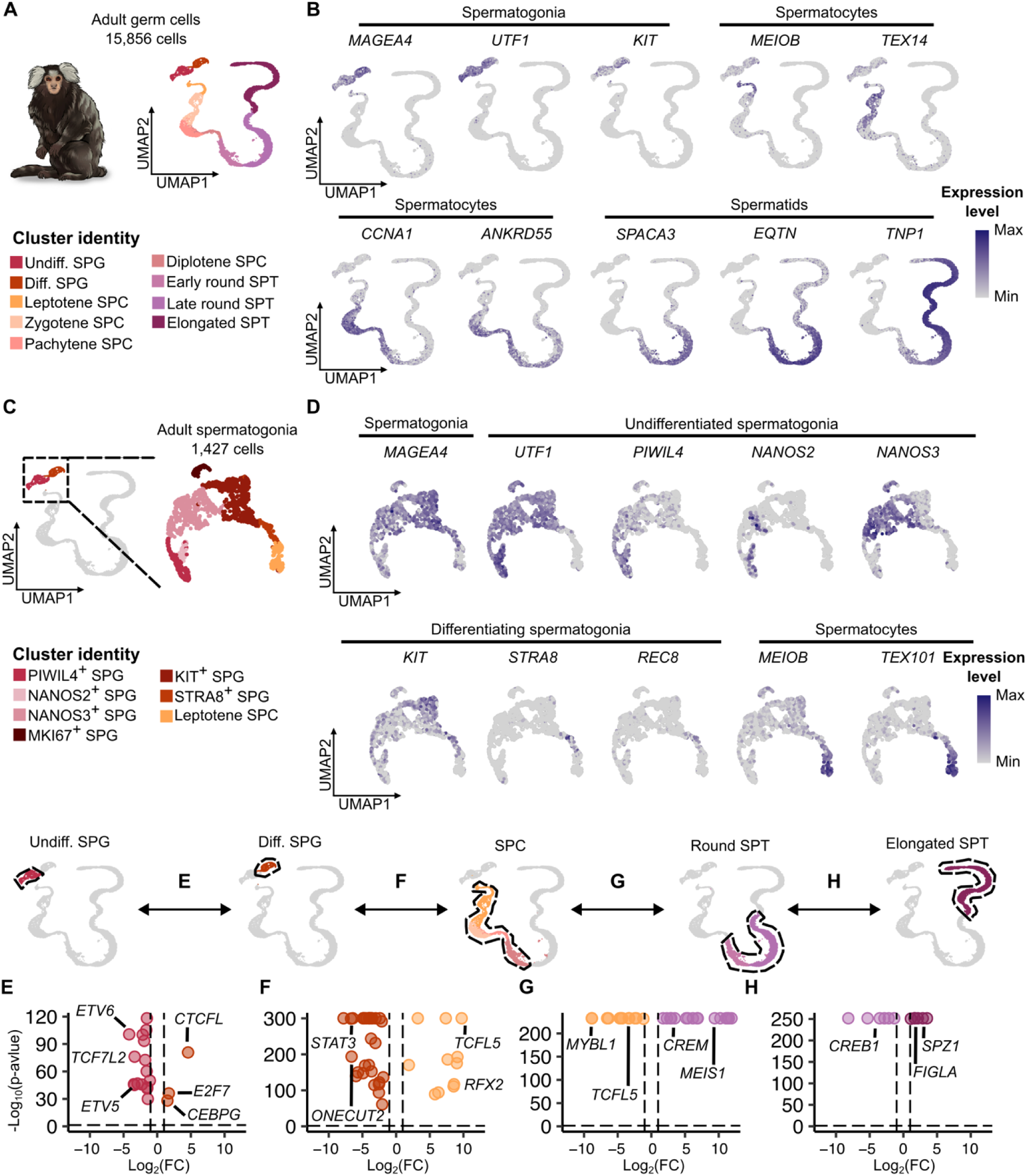
Regulon activity governing germ cell differentiation from spermatogonia to sperm. A) UMAP plot of the adult germ cell subset (15,856 cells). DEGs are provided in **Table S5**. B) Feature plots showing the genes used for cell assignment. C) UMAP plot of the adult spermatogonial subset (1,427 cells). DEGs are provided in Table S5. D) Feature plots showing the genes used for cell assignment. E-H) Results of the differential regulon activity analysis. UMAP plots (top row) highlight the clusters considered for each comparison and volcano plots (bottom row) depict respective results. Vertical dashed lines represent the fold change threshold (|log_2_(fold change) = 1|), and the horizontal dashed line represents the −log_10_(p-value) threshold (p-value = 0.05). Differentially active regulons are provided in **Table S5**.

To scrutinize the composition of the adult spermatogonial compartment, we subset and re-clustered the undifferentiated and differentiating spermatogonia consisting of 1,428 cells (**Fig. 5C**; **Table S5**). We identified the same spermatogonial makeup (**Fig. 5C** and **5D**) as for the pubertal stage (**Fig. 4A** and **4B**). Specifically, *PIWIL4*^+^, *NANOS2*^+^, and *NANOS3*^+^ spermatogonia formed the undifferentiated spermatogonial compartment, a subpopulation of *NANOS3*^+^ spermatogonia showed a proliferative profile (*MKI67*) (**Fig. S5A** and **S5B**), and *KIT*^+^, *STRA8*^+^ and *REC8^+^*spermatogonia represented the differentiating spermatogonia, linking to spermatocytes (*MEIOB*, *TEX101*) (**Fig. 5C** and **5D**). Moreover, differential regulon analysis of the adult spermatogonial subset revealed overlapping regulons between the same cell identities throughout development. For instance, *SOX4* activity was inferred in every undifferentiated spermatogonial clusters (*PIWIL4*^+^, *NANOS2*^+^, and *NANOS3*^+^), *OCT6* for *PIWIL4*^+^ spermatogonia, and *ETV6* for *NANOS3*^+^ spermatogonia (**Fig. S5C**; **Table S5**).

Seeking to identify the core transcription factors that define the identities of the adult germ cell types, we compared regulon activity among undifferentiated spermatogonia (*PIWIL4*^+^, *NANOS2*^+^, and *NANOS3*^+^), differentiating spermatogonia (*KIT*^+^ and *STRA8*^+^), spermatocytes (leptotene, pachytene, zygotene, and diplotene), round spermatids (early and late round), and elongated spermatids (**Fig. 5E-H**; **Table S5**). Among the regulons active in undifferentiated spermatogonia were *ETV5*, *ETV6*, and *TCF7L2* (**Fig. 5E**; **Table S5**), which are associated with stem cell self-renewal.^34–36^ Differentiating spermatogonia showed differential regulon activity of *CTCFL*, *CEBPG,* and *E2F7* when compared to undifferentiated spermatogonia (**Fig. 5E**; **Table S5**) and *STAT3* and *ONECUT2* in comparison to spermatocytes (**Fig. 5F**; **Table S5**). While CEBPG has been shown to promote cell cycle progression,^37^ the E2F7-dependent transcriptional program has been shown to regulate cell cycle protecting genome integrity^38^ and to orchestrate differentiation of multiciliated cells^39^, ascertaining our findings in highly proliferative differentiating spermatogonia (**Fig. S5A**). Regulons active in spermatocytes compared to differentiating spermatogonia included *TCFL5*, whose deficiency leads to meiotic arrest in mice,^40^ and *RFX2*, a major transcriptional regulator for spermiogenesis.^41^ When compared to round spermatids, *MYBL1* and *TCFL5* regulons were differentially active (**Fig. 5G**; **Table S5**). The transcriptional profile of round spermatids included genes regulated by *MEIS1* and *CREM* when compared to spermatocytes (**Fig. 5G**; **Table S5**), and *CREB1* when compared to elongated spermatids (**Fig. 5H**; **Table S5**). *CREM* is a well-known gene that is essential for spermiogenesis^42^ while *CREB1* is predicted to upregulate *CATSPER1*, essential for fertilization.^43^ Elongated spermatids showed increased activity of the regulons controlled by *SPZ1* and *FIGLA* when compared to round spermatids (**Fig. 5H**; **Table S5**), with SPZ1 being also indispensable for correct spermiogenesis.^44^

In summary, we uncovered that the makeup of the adult spermatogonial compartment is the same as at the onset of puberty. Moreover, we identified the molecular gatekeepers involved in meiosis (*TCFL5*, *MYBL1*) and spermiogenesis (*RFX2*, *CREM*), paving the way for full male germ cell differentiation.

### The molecular fingerprint of spermatogonial subpopulations throughout development

Finally, we set out to assess if the different spermatogonial subpopulations maintain their molecular features throughout postnatal development. Undifferentiated spermatogonia represented 38.4% of the neonatal, 100% of the pre-pubertal, 59.6% of the pubertal, and 4.9% of the adult germ cells among the scRNA-seq datasets (**Fig. 6A**; **Table S2-S5**). Interestingly, the proportions of the spermatogonial subpopulations captured in pubertal and adult tissues, in particular of *PIWIL4^+^*, *NANOS2^+^,* and *NANOS3*^+^ spermatogonia, remained similar (**Fig. 6B**; **Table S6**).

**Figure 6:**
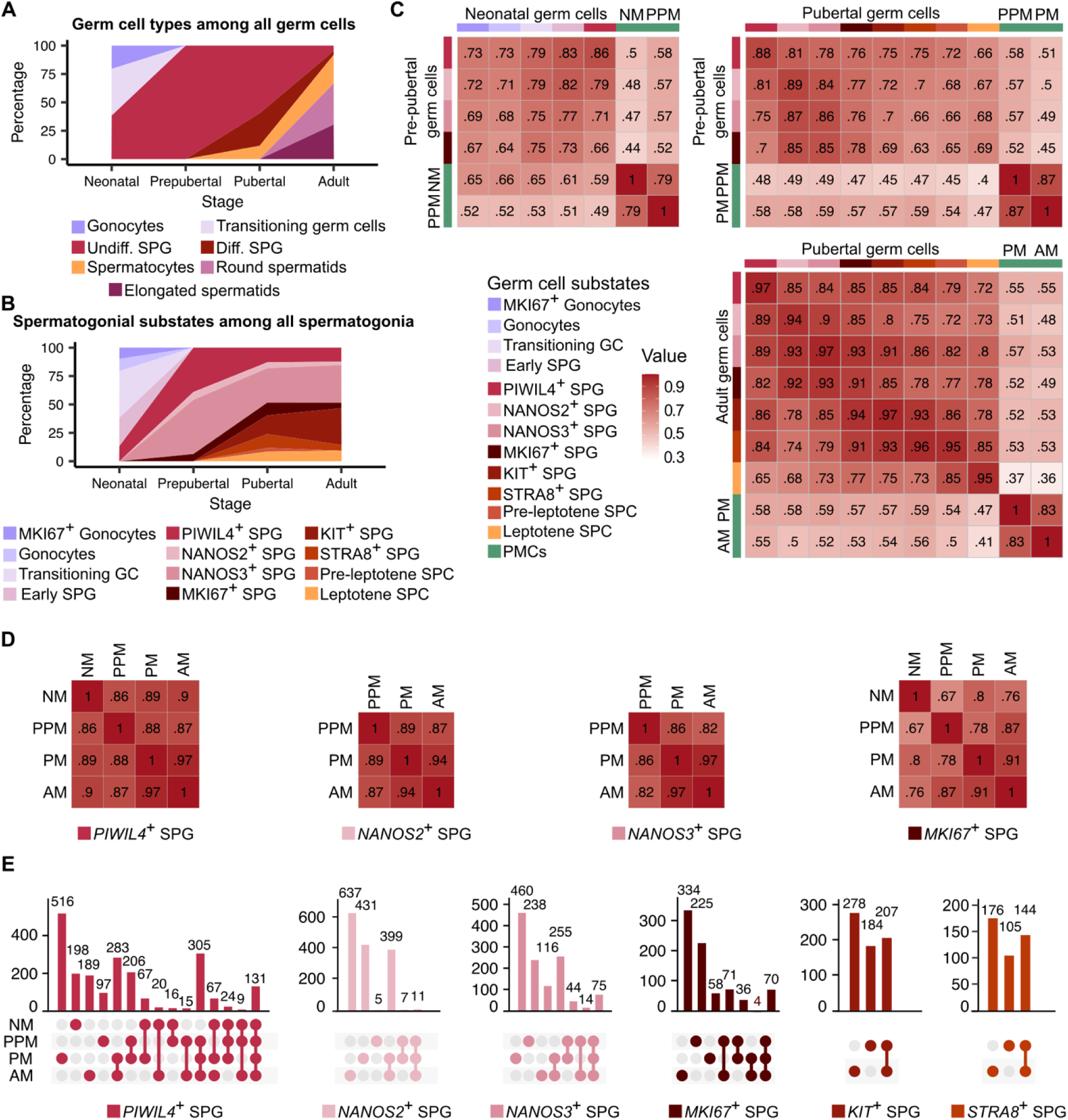
The molecular fingerprint of spermatogonial subpopulations throughout development. A) Area plot portraying the changes in germ cell type proportions during development. B) Area plot depicting changes in spermatogonial state proportions during development. C) Heatmap showing the results of a Pearsońs correlation between transcriptomes of each germ cell/spermatogonial cluster identity, grouped by developmental stage comparing neonatal (NM) vs. pre-pubertal (PPM), pre-pubertal vs. pubertal (PM), and pubertal vs. adult (AM). Results for peritubular myoid cells (PMCs) are displayed for comparison. D) Heatmaps showing Pearsońs correlation results, according to spermatogonial cluster identity, at every developmental stage. E) UpSet plots depicting the overlap of the top differentially expressed genes (DEGs; up to 500) between the developmental stages in spermatogonial substates. Full lists of overlapping genes are provided in **Table S6.**

To scrutinize whether the transcriptional states of spermatogonia at the four developmental stages shared a common molecular fingerprint, we performed Pearson’s correlations of their transcriptomes. Importantly, the strongest correlations were among those cells with the same cell identity. For instance, comparing neonatal and pre-pubertal *PIWIL4*+ spermatogonia resulted in the strongest significance (*r* = 0.86; **Fig. 6C**), as was the case for the comparison between pubertal and adult stages (*r* = 0.97) (**Fig. 6D**), highlighting how little the transcriptome of each spermatogonial substate changes during development. Similarly strong correlations were observed for the *NANOS2^+^* and *NANOS3*^+^ subpopulations (**Fig. 6D**), detectable from the pre-pubertal stage onwards, indicating that, once the different spermatogonial subtypes are established, their transcriptional fingerprint is maintained throughout life. To explore the spermatogonial transcriptional fingerprint, we identified marker genes shared by each spermatogonial subpopulation during development by overlapping their DEGs lists (**Table S6**). We uncovered that *PIWIL4*^+^ spermatogonia shared 131 DEGs in each developmental stage (*e.g. SPOCD1, C22H19orf84*), *NANOS2^+^* spermatogonia 11 (e.g. *FGD3, IFIT3)*, and *NANOS3*^+^ spermatogonia a total of 75 shared DEGs (e.g. *DPPA4, CITED2*) (**Fig. 6E**; **Table S6**). These marker genes therefore represent the molecular identity of the distinct spermatogonial subpopulations.

To follow the dynamics of the undifferentiated spermatogonial compartment at protein level, we performed quantitative histomorphometrical analyses (**Fig. S6A**). The number of MAGEA4^+^ spermatogonia increased continuously and significantly from the neonatal to the adult stage (neonatal: 2.5; adult: 19.7 MAGEA4^+^ cells/RT) (**Fig. S6B**). While the same general pattern was found for the UTF1^+^ spermatogonia, this increase was more subtle (neonatal: 1.6; adult: 4.8 UTF1^+^ cells/RT) (**Fig. S6B**). Importantly, the number of spermatogonia positive for PIWIL4 remained stable throughout development (neonatal: 3.2; adult: 2.9 PIWIL4^+^ cells/RT) (**Fig. S6B**). Interestingly, NANOS3^+^ cells showed a similar pattern with slightly increased numbers in pre- and pubertal tissues (neonatal: 1.6; pre-pubertal: 2.6; pubertal: 2.4; adult: 0.9 NANOS3^+^ cells/RT) (**Fig. S6B**; **Table S6**).

Finally, we assessed the ability of the marker-based spermatogonial subpopulations to form reserve A_dark_ spermatogonia, cells characterized by highly condensed nuclear chromatin, low proliferative activity, and the capacity to repopulate the testis after injury.^45,46^ We identified A_dark_ spermatogonia from pre-pubertal stage onwards (**Fig. S6C**). While 9.7% of the adult MAGEA4^+^ cell population was A_dark_ spermatogonia, UTF1 expression was found in the majority of A_dark_ spermatogonia (>70%) (**Fig. S6C**), which is comparable to what is found in human testes.^5^ Interestingly, PIWIL4^+^ and NANOS3^+^ A_dark_ spermatogonia were found in higher numbers in pre-pubertal tissues (41.3% and 27.3%, respectively) but decreased towards the adult stage (13.3% and 9.3%, respectively) (**Fig. S6C**), aligning with the enlargement of the seminiferous tubules and increase in meiotic and post-meiotic germ cell types. These datasets highlight the general ability of all spermatogonial subpopulations to form A_dark_ reserve cells, however at different capacities. Interestingly, and in contrast to the other marker-based subpopulations of undifferentiated spermatogonia, very few DPPA4^+^ A_dark_ spermatogonia were identified during development (pre-pubertal: 3.3%; pubertal: 2%; adult: 6.7%; **Fig. S6D; Table S6**). The fact that we found few DPPA4^+^ A_dark_ spermatogonia in general either hints at a role of DPPA4 in regulating spermatogonial chromatin structure or the requirement of an A_pale_ morphology for expression of DPPA4.

To conclude, numbers of undifferentiated spermatogonial subpopulations (*PIWIL4*^+^, DPPA4^+^) remain stable throughout postnatal development, as does their transcriptional fingerprint and molecular identity, ensuring the perpetuity of the male germ line.

## Discussion

To date, the regulatory networks shaping the spermatogonial compartment and directing germ cell differentiation throughout human development have remained largely unknown. These knowledge gaps hinder advancement of research on testicular germ cell cancers, caused by persisting gonocytes,^47^ preclude treatment of male infertility,^11^ and development of fertility restoration approaches.^12^ To close these gaps, we used the marmoset as the best available non-human primate model to scrutinize the molecular makeup of the spermatogonial compartment throughout development and to uncover the gatekeepers of germ cell differentiation.

Throughout development, we uncovered both distinct and common cell-type characteristics. Unique to the neonatal stage was the transition from gonocytes to *PIWIL4*^+^ spermatogonia, which occurs via two linking cell clusters (**Fig. 7A**). Among the transcriptional gatekeepers governing this transition was CITED2, a transcriptional co-factor that interacts with AP factors such as AP2γ to regulate gene expression, and also competes with hypoxia-induced factors for their binding sites in target gene promoters, downregulating them.^48^ The latter may be especially relevant for the metabolic changes during the migration towards the tubular walls in the gonocyte-to-spermatogonia transition.^49^ Also, CITED2 is involved in the suppression of pluripotency markers (e.g. OCT4) in humans,^50^ and aberrant expression has been linked to several cancers, such as breast, colon, and prostate cancer.^51,52^ Therefore, we hypothesize that CITED2 plays a major role in the gonocyte-to-spermatogonia transition and is a candidate gene for germ cell tumorigenesis.

**Figure 7:**
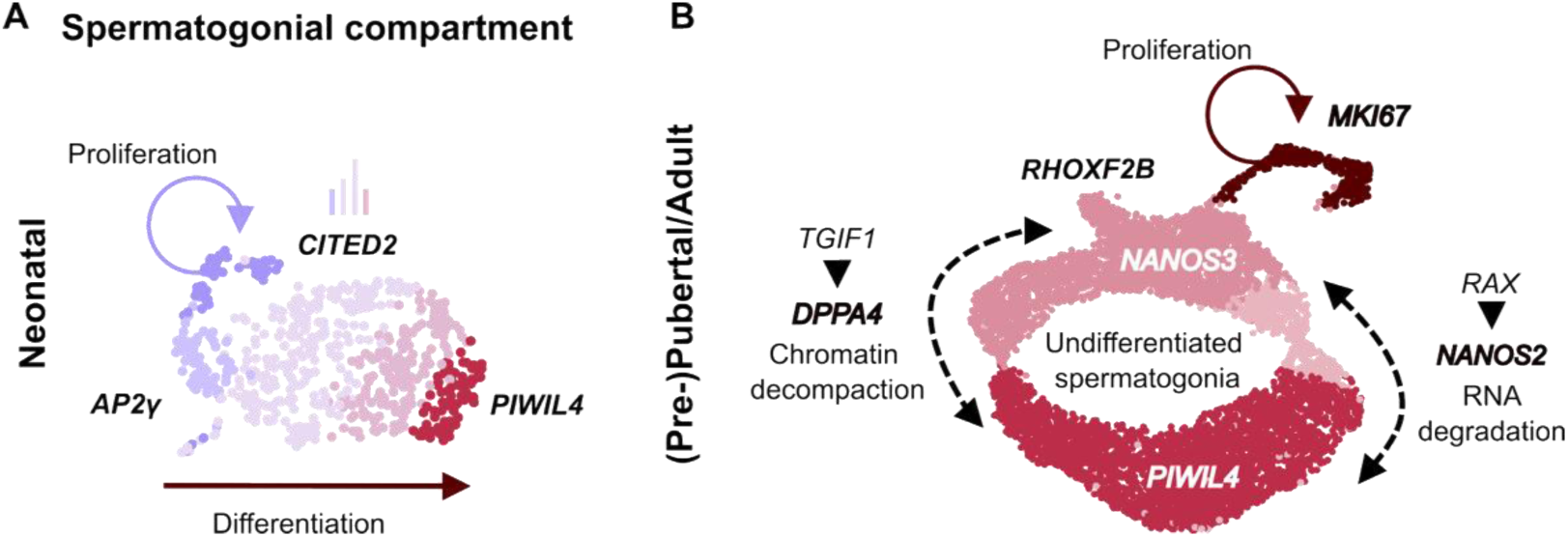
Model of cellular states and dynamics in the spermatogonial compartment throughout development. A) Model of the gonocyte-to-spermatogonia transition. B) Model of the spermatogonial compartment plasticity and differentiation.

Consistent throughout development was the presence of PIWIL4^+^ spermatogonia, with a remarkably constant transcriptional profile. The detection of these cells in neonatal marmoset testes seems to be in contrast to a previous study, which detected PIWIL4^+^ spermatogonia in marmoset only from three months of age onwards.^53^ Our findings however align with data from macaques and humans, in which PIWIL4^+^ spermatogonia were detected in fetal testicular tissues.^54,55^ Functionally, and based on mouse data, PIWIL4 is indispensable for maintaining DNA integrity by silencing transposable elements via *de novo* DNA methylation in gonocytes.^7,8^ Importantly however, the timing of *de novo* DNA methylation in the marmoset is different from that in mice, with increases of global DNA methylation continuing beyond the pubertal stage. ^56^ Moreover, in rodents and humans, global methylation levels change significantly during each wave of spermatogenesis, a period when silencing of transposable elements (TE) is of critical importance.^57,58^

Regarding the organization of the spermatogonial compartment, from pre-pubertal stage onwards, similar to the human situation, *PIWIL4*^+^ and *NANOS3*^+^ spermatogonia form a roundabout continuum of cells (**Fig. 7B**).^3,5^ We found that *NANOS2* and *DPPA4,* regulated by *RAX* and *TGIF1*, respectively, are crucially relevant for the two transition paths. NANOS2 is a highly conserved, testis-specific RNA-binding protein important for fetal development and post-natal spermatogonial stem cells self-renewal that acts by regulating essential transcripts for germ cell determination via deadenylation.^59^ DPPA4 primes chromatin through regulation of H3K4me3 and H3K27me3 at bivalent domains, maintaining developmental competency.^60^ This activity is related to global DNA decompaction, which aligns with the scarcity of DPPA4^+^ cells showing an A_dark_ morphology.^61^ We hypothesize that *PIWIL4*^+^ spermatogonia give rise to *NANOS3*^+^ spermatogonia, which undergo asymmetrical division requiring silencing of transposable elements in one of the cells, while the other becomes primed for differentiation, as suggested for the rhesus monkey and the human.^62,63^ As differentiation only occurs upon puberty, this may explain the peak in numbers of NANOS3^+^ spermatogonia in pre-pubertal tissues. Alternatively, NANOS2 and DPPA4 may constitute two alternative transitioning paths into NANOS3^+^ cells, ensuring robustness of the system as described for megakaryopoiesis.^64^

The population of proliferatively active germ cells is distinct between the developmental time points. While AP2γ ^+^ gonocytes showed proliferative activity in neonatal tissues, NANOS3^+^ cells form the hub of spermatogonial expansion from pre-puberty onwards, based on MKI67 expression. From the pubertal stage onwards, this is complemented by proliferative activity among differentiating *KIT*^+^ cells, facilitating expansion of both compartments, in line with data from the adult rhesus monkey.^62^ These findings may explain the distinct impact of gonadotoxic treatments administered at different developmental stages on long-term fertility.^65^

Comparison of the pre-pubertal and pubertal spermatogonial compartments showed that *RHOXF2B* upregulation marked the origin of differentiation out of the *NANOS3*^+^ spermatogonia (**Fig. 7B**). In line with this, RHOXF2B is also expressed by differentiating B spermatogonia^66^ in human adult testicular tissues. Reproductive homeobox factors have been associated with TE repression in the murine germline^67^ and mutations in the *RHOXF2* gene were associated with infertility in humans, likely due to uncontrolled LINE1 expression in spermatogonia,^68^ stressing the relevance of TE silencing in the germline.

In conclusion, we report that the spermatogonial stem cell system is initially formed via upregulation of CITED2. Moreover, we identified NANOS2 and DPPA4 as regulators of spermatogonial plasticity and uncovered the molecular regulators of male germ cell differentiation during postnatal marmoset development, which will inform research on testicular germ cell tumors, male infertility, and fertility restoration approaches.

### Limitations

In this study, we utilized marmoset testicular tissues spanning different developmental time points to study transcriptional regulation during primate male germ cell differentiation. While these experiments were not performed in humans, marmoset monkeys have been extensively used as an accurate model for the study of human postnatal germ cell differentiation. Despite the high number of cells analyzed at each step of this article, the number of samples for scRNA-seq (n=11) and histomorphometrical analyses (n=24) were limited for ethical and legal reasons.

## Resource availability

### Lead contact

Further information and requests for resources and reagents should be directed to, and will be fulfilled by the lead contact, Nina Neuhaus.

### Materials availability

This study did not generate new unique reagents.

### Data and code availability

scRNA-seq data are deposited in ArrayExpress with the accession number E-MTAB-16798. This paper does not report original code.

Any additional information required to reanalyze the data reported in this paper is available from the lead contact upon request.

## Supporting information

Key Resource Table

## Acknowledgements

We thank Nicole Terwort, Reinhild Sandhowe, Heidi Kersebom, Elke Kößer, Vesna Bojovic, and Jutta Salzig for their excellent technical support, Lidia Jaen Muñoz for the marmoset illustrations, the CeRA for institutional funding, and the invaluable support of the Central Animal Facility of the Faculty of Medicine animal caretakers.

This work was funded by the Innovative Medizinische Forschung (IMF) of the Medical Faculty of the University Münster to N.N. and S.L. (LA112010) and the German Federal Ministry of Research, Technology and Space (BMFTR) as part of the project ReproTrack.MS – Centre for Research and Development of Reproductive Scientists (01GR2303), and with a grant to G.M.z.H. as part of the ‘Lipid Immune Neuropathy Consortium’. Moreover, this work was funded by the German Research Foundation (DFG) with grants to N.N. and S.L. (LA 4064/6-1 | NE 2190/7-1; LA 4064/4-1), A.S.B (464240267), and by grants to G.M.z.H. (ME4050/12-1, ME4050/13-1, ME4050/18-1 and a grant in the Interdisciplinary Koselleck programme).

## Author contributions

Conceptualization, J.M.P., S.D.P., S.L., and N.N.; Methodology, J.M.P, S.D.P., M.W., T.L., A.O., and J.W.; Software, J.M.P., M.W., T.L., Sa.S.; Investigation, J.M.P., S.D.P., M.W., T.L., S.L., and N.N.; Resources, X.L., G.M.Z.H., JW, and St.S.; Writing – Original Draft, J.M.P., S.L., and N.N.; Writing – Review & Editing, all authors; Visualization, J.M.P.; Funding Acquisition, S.L. and N.N.; Supervision, S.L. and N.N.

## Supplemental information titles and legends

Document S1. Figures S1–S6.

**Table S1. Supplemental data of scRNA-seq analyses and histological quantifications, related to Figure 1**.

(A) Samples included in the study.

(B) Cell numbers after filtering steps.

(C) Cell numbers per testicular cell type per developmental stage.

(D-H) DEGs of neonatal (D), pre-pubertal (E), pubertal (F), adult (G) testicular cells and neonatal immune cells (H).

(I-L) GO terms of neonatal (I), pre-pubertal (J), pubertal (K), and adult (L) testicular cells.

(M) Overlap of GO terms.

(N) Histological quantifications.

**Table S2-S5. Supplemental data of scRNA-seq analyses of neonatal (S2), pre-pubertal (S3), pubertal (S4), and adult (S5) germ cells and histological quantifications, related to Figures 2**, **3**, **4 and 5, respectively.**

(A) Number of germ cells per cluster and cell cycle analysis.

(B) Germ cell-specific differentially expressed genes.

(C) Functional annotation (GO terms) of germ cells per cluster.

(D) Gene regulatory networks in the germ cells per cluster.

(E) Sequential regulon activity analysis results in the germ cells.

(F) List of regulons and their predicted regulated genesets.

(G) Histological quantifications.

**Table S6. Supplemental data on the molecular fingerprint of spermatogonial subpopulations, related to Figures 6 and S6.**

(A) Pearsońs correlations between spermatogonial identities per developmental stage and per cluster.

(B) Overlap of DEGs between developmental stages per spermatogonial identity.

(C) Histological quantifications.

## Materials and methods section

### EXPERIMENTAL MODEL AND STUDY PARTICIPANT DETAILS

#### Marmoset monkeys (*Callithrix jacchus*)

Testicular tissues from neonatal (n=2; 0-2 days old), pre-pubertal (n=3; 6 months old), pubertal (n=2; 8 months old), and adult male marmosets (n=4; >1 years old) were obtained from the colony available at the Central Animal Facility (ZTE) of the University Hospital of Münster. Marmosets were kept in pairs/families under a 12 h light/12 h darkness regimen, with identical exercise conditions and unlimited access to tap water. Animals were fed food pellets from Altromin (Lage, Germany) composed for marmosets, and food was supplemented daily with fresh fruits and vegetables. Tissues from male marmoset monkeys were obtained in accordance with the German Animal Protection Law under the animal licence No. LANUV AZ 81-02.05.05.21.014. Additional blocks of paraffin-embedded fixed testicular tissues stored at the CeRA (n=11) were included for cohort completion (Supplemental Table 1).

### METHOD DETAILS

#### Histological tissue preparation

Pieces of testicular tissue samples were fixed overnight in Bouin’s solution or paraformaldehyde (PFA; 4%) and dehydrated in 70% ethanol or increasing concentrations of ethanol (30%, 50%, 70%), respectively. The tissue was paraffin-embedded, cut into 5 μm-thick sections, and mounted onto Superfrost Plus Adhesion Microscope Slides (Epredia, Cat# 4951PLUS-001).

#### Stainings

##### Periodic acid-Schiff/hematoxylin staining

Bouin-fixed sections were deparaffinized in AppiClear (Applichem, Cat# A4632.2500), rehydrated in decreasing ethanol concentrations (99%, 96%, 80%, 70%), and washed in distilled water. The slides were sequentially incubated with periodic acid (Sigma-Aldrich, Cat# 1.00524.0100), Schiff’s reagent (Sigma-Aldrich, Cat# 1.09033.0500), and counterstained with Mayeŕs hematoxylin (Merck, Cat# 1.09249.0500). Samples were dehydrated in increasing ethanol concentrations (70%, 80%, 96%, 99%) and mounted with Merckoglass (Merck, 1.03973.0001) under a glass coverslip.

##### Immunohistochemical staining

Immunohistochemical stainings were performed against DAZL, OCT4, AP2γ, LIN28A, MAGEA4, PIWIL4, UTF1, NANOS3, and DPPA4. The tissue sections were deparaffinized in Appiclear and rehydrated as described above. Heat-induced antigen retrieval was performed using sodium citrate buffer (pH 6.0) up to boiling point using a microwave (1x5 min at 800W, 2x5 min at 400W). The samples were blocked with bovine serum albumin (BSA, 5%; Sigma-Aldrich, Cat# A9647) and goat serum (Sigma-Aldrich, Cat# G6767), followed by incubation with primary antibodies diluted in blocking solution (See STAR Methods table) at 4°C overnight. Sections were incubated for 1 hour with the respective biotin-conjugated secondary antibodies against mouse or rabbit IgG (See STAR Methods table). Sections were washed and incubated with streptavidin-horseradish-peroxidase (Sigma-Aldrich, Cat# S5512) and 3,30-diaminobenzidine tetrahydrochloride solution (Applichem, Cat# A0596.0001) as substrate. The reaction was stopped with distilled water. Prior to dehydration in increasing ethanol concentrations and clearing in Appiclear, nuclei were counterstained with Mayer’s hematoxylin. The sections were mounted with Merckoglass under glass coverslips and scanned using a Precipoint M8 Microscope and Scanner (Precipoint, Freising, Germany).

##### Immunofluorescence stainings

The following immunofluorescence co-stainings were performed: LIN28A and MAGEA4, AP2γ and MKI67, NANOS3 and MKI67, EGR1 and MAGEA4, AP2γ, CITED2 and PIWIL4, and NANOS3, DPPA4 and PIWIL4. The tissues were deparaffinized in Appiclear, rehydrated in decreasing ethanol concentrations (99%, 96%, 80%, and 70%), and washed with distilled water. Antigen retrieval was performed as described above. Autofluorescence was quenched with 1 M Glycine (Sigma-Aldrich, Cat# G7126) and tissues were permeabilized with 0.1% Triton X-100 (Sigma-Aldrich, Cat# 93443) in Tris-buffered saline (TBS) for 15 minutes in a wet chamber at room temperature. Unspecific antibody binding was blocked with donkey serum (5%; Sigma-Aldrich, Cat# S30), BSA (1%) and Tween 20 (0.1%v/v; Sigma-Aldrich, Cat# 655205) in TBS for 30 minutes before incubation overnight with combinations of the primary antibodies (See STAR Methods table). The tissue sections were washed and incubated with species-specific secondary antibodies conjugated with different fluorophores for 1 hour at room temperature and with the conjugated secondary antibodies for 1 hour (See STAR Methods table). In case of primary antibodies in the same species (i.e. AP2γ, CITED2 and PIWIL4, or NANOS3, DPPA4 and PIWIL4 co-stainings), FlexAble 2.0 CoraLite Plus kits (Proteintech, Cat# KFA501, Cat# KFA502, and Cat# KFA503) were used to conjugate the primary antibodies to different fluorophores following the supplier’s protocol. In this case, the tissues were incubated with the conjugated primary antibodies for 2 hours at room temperature. The slides were mounted under glass coverslips with VectaShield Mounting Medium containing 4,6-diamidino-2-phenylindole as nuclear counterstain (Vector Laboratories, Cat# H-1200). Immunological signals were captured using an Olympus BX61VS microscope or Evident Slideview VS200 Universal whole slide imaging scanner and evaluated using the software VS-ASW-S6 (Olympus, Hamburg, Germany).

#### Histological quantifications

##### Point Counting

The composition of seminiferous tubules, interstitium, and blood vessels was studied using a point counting approach based on ten non-overlapping fields of the PAS-stained sections per animal (n=12) obtained at 60x magnification (area ≍ 1 mm^2^) with the Precipoint M8 Microscope and Scanner (Precipoint, Freising, Germany). Briefly, 50 points randomly scattered through each snapshot (total of 500 points) were assigned by an observer to the different categories. Points that could not be assigned to a category were recorded as non-assigned (NA). The proportions were calculated over the total of assigned points and multiplied by 100.

##### Most advanced germ cell stage

For the assessment of the most advanced germ cell type identifiable in seminiferous tubules, we used MAGEA4-stained tissues of marmoset (n=12). Scans were obtained using a Precipoint M8 Microscope and Scanner (Precipoint, Freising, Germany) and quantifications were performed using the Viewpoint software (Precipoint, Freising, Germany). We categorised a total of 100 seminiferous tubules per animal as showing a Sertoli cell-only phenotype or with spermatogonia, spermatocytes, round spermatids, or elongated spermatids as most advanced germ cell type present.

##### Germ cell markers per round tubule

Positive cells for each marker (DAZL, OCT4, AP2γ, LIN28A, MAGEA4, UTF1, PIWIL4, NANOS3, DPPA4) were quantified in 20-40 (depending on tissue size) round tubules randomly selected across each tissue section per animal. Tubules were considered as round if the ratio between the shortest and longest diameters was 1.0-1.5. In case the minimum number of round tubules could not be achieved in a single section, an independent section, at least 15 μm away from the first one, was additionally analysed. Quantifications were performed using the Viewpoint software for Precipoint scans.

##### Co-labelling index in IF-stained tissue sections

A total of 200 MAGEA4^+^, 100 NANOS3^+^, or 50 (when possible) AP2γ^+^ cells were quantified using individual immunofluorescence co-stainings. The percentage of MKI67^+^ among the NANOS3^+^ cells, and of EGR1^+^ among MAGEA4^+^ cells was calculated in neonatal, pre-pubertal, pubertal, and adult PFA-fixed marmoset testicular tissue sections (n=3 each stage, except for PM: n=2).

##### Co-localization analysis in testicular tissue sections

In the cases of AP2γ, CITED2 and PIWIL4, and LIN28A and MAGEA4, random tubules were selected using DAPI signal, and the number of single, double, and triple positive cells in the case of AP2γ, CITED2, and PIWIL4, were quantified until reaching a total of 200 cells. The percentage of cells expressing each marker combination was calculated.

##### Localization of neonatal germ cells

Positive cells for AP2γ, LIN28A, MAGEA4, and PIWIL4 were categorized into luminal or basal based on their location within the seminiferous tubules of neonatal marmoset testicular tissues (n=7). Cells showing protrusions towards the peritubular wall were categorized as basal. 50 positive cells were quantified for each marker, when possible. If not, as it was the case for individual samples, all positive cells contained in one tissue section were considered (**Table S2**).

#### Single-cell RNA-sequencing

##### Testicular tissue digestion

Single-cell suspensions were obtained from marmoset testis using a two-step enzymatic tissue digestion. Briefly, following removal of the tunica albuginea, testis tissues were cut into 1x1 mm^3^ pieces. For the first enzymatic digestion step, tissues were incubated in Minimum Essential Medium Alpha (MEMα; Life Technologies GmbH, Gibco, Cat# 22561-021) with 1 mg/ml collagenase (Sigma-Aldrich, Cat# C9891) for 10 minutes and inversion every 3 minutes. This reaction was stopped by addition of MEMα supplemented with 10% fetal bovine serum (Sigma-Aldrich, Cat# S0615) and 1% penicillin and streptomycin (Life Technologies GmbH, Gibco, Cat# 15140-122). Subsequently, the tissue fragments were incubated with trypsin (Life Technologies GmbH, Gibco, Cat# 25725-018) and DNAse (Sigma-Aldrich, Cat# SLCJ6659) [4 mg/ml and 2.2 mg/ml, respectively, in Hank’s Balanced Salt Solution (Life Technologies GmbH, Gibco, Cat# 14175-053)]. Due to differences in tissue composition, neonatal tissues were exposed to lower concentrations of trypsin (2 mg/ml). A hemolysis buffer [(8.29 g/l NH_4_Cl (Merck, Cat# 1145), 1 g/l KHCO_3_ (Sigma, Cat# P-9144), 0.037 g/l EDTA (Serva, Cat# 11278) in bi-distilled water] was used to remove red blood cells. The cells were washed in MEMα supplemented with 10% fetal bovine serum with 1% penicillin and streptomycin three times before passing through a 70 µm strainer, obtaining a single-cell suspension.

##### Spermatid depletion by density gradient

A density gradient was used to deplete haploid cells and enrich diploid and double-diploid cells in two adult testicular single-cell suspensions. 20 ml of 4% BSA in MEMα and 2% BSA in HBSS were overlaid one on top of the other, allowing the visualization of an interface between the two solutions. 1 ml of 0.5% BSA in MEMα was dispensed over the 2% BSA fraction. The cells were suspended in 1 ml of the 0.5% BSA in MEMα and pipetted carefully over the top fraction, with the same BSA concentration. After 1 hour of sedimentation, 2 ml aliquots were eluted from the bottom of the column with a flow of 2ml/min and collected in separate tubes. The number of live cells per fraction was quantified using a Neubauer chamber and trypan blue staining, confirming integrity of the cells after the sedimentation gradient.

To confirm the depletion of haploid cells, we performed ploidy analysis in 100 µl of each fraction. After centrifugation, the cells were incubated in the dark for 30 min with a solution containing 50 mg/ml Propidium Iodide (Sigma- Aldrich, Cat# P4170), 1 mg/ml bovine serum albumin (Sigma-Aldrich, Cat# A9647), 0.1% Triton X-100 (Sigma-Aldrich, Cat# 93443), and 10 mg/ml RNase A (Sigma-Aldrich, Cat# R6513) in phosphate-buffered saline (PBS). After incubation, approximately 10,000 cells were analysed for each sample using a Beckman Coulter CytExpert QC Flow Cytometer (Beckman Coulter, Krefeld, Germany/Brea, California). Debris was defined based on forward and side scatter and excluded from the analysis. The cells analysed for DNA content were measured using a laser at 617 nm and assigned to the following categories according to staining intensity: 1) haploid cells (spermatids and sperm), 2) diploid cells (spermatogonia and somatic cells, including Leydig, peritubular myoid, and Sertoli cells), 3) double-diploid cells (primary spermatocytes). Fractions 0-10 contained mostly diploid and double-diploid cells, so they were pooled together and subjected to scRNA-seq.

##### ScRNA-seq library preparation and sequencing

12,000 cells per marmoset sample (neonatal = 2; pre-pubertal = 3; pubertal = 2; adult = 4) were suspended at a concentration of 500 cells/ml in MEMα and loaded into the Chromium Single Cell Chip. Library preparation was performed following the kit instructions (Chromium Single Cell Kit v2 chemistry for adult marmosets 1 and 2 and v3 chemistry for all the other samples). Briefly, ∼6,000 cells per sample were captured by the 10x Genomics Chromium controller, cDNA was synthesized, and 12-14 cycles were used for library amplification according to the 10x protocol. The resulting libraries were quantified using a Lab901 TapeStation system (Agilent, Santa Clara, United States) before shallow paired-end sequencing (NextSeq 550 sequencer; Illumina, San Diego, United States) for quality control. The final, deep sequencing was performed on a NovaSeq 6000 sequencer (Illumina, San Diego, United States), using 2 x 150 bp paired-end sequencing.

##### Reference genome and read count assignment

We used a modified version of the Callithrix_jacchus_cj1700_1.1 reference (RefSeq assembly accession GCF_009663435.1) for downstream analyses. A CellRanger (v3.1.0) reference was created with the modified Callithrix_jacchus_cj1700_1.1 reference using its mkref command. Gene counts were computed by CellRanger’s (v6.0.2) count command for each of the 11 samples using default parameters and including intronic reads. Although this reference contains both mitochondrial sequences and gene locations, location information was flagged exclusively as ‘CDS’ (CDS, coding sequence) in the references’ GTF file. This prevented assignment of gene counts to mitochondrial genes, which are required for downstream quality control. To work around this issue, ‘CDS’ flags in mitochondrial gene annotations were replaced with ‘exon’ flags, thus enabling gene counts to be assigned to mitochondrial genes.

##### Quality control for single-cell RNA-seq data

CellRanger counts were imported using the Seurat toolbox (v5.0.3),^69^ resulting in a total of 68,293 imported cells before quality control (QC) filters. SoupX^71^ was applied, which identifies the ambient RNA transcripts based on distribution throughout the different clusters, calculates a coefficient of contamination, and removes the ambient RNA reads from the expression matrices, enhancing clustering. Cells were filtered based on the total number of genes detected and percentage of counts assigned to mitochondrial genes. Due to variable count distributions among samples and Seurat clusters, we selected the lower threshold per individual. Cells were filtered out if the total number of detected genes in that cell was <500 in NM3, <800 in all PPM samples and PM2, <700 in PM3, AM1 and AM12, <600 in AM2, and <1,000 in AM13. No threshold in the nFeature_RNA was selected for NM2 due to a lower sequencing depth compared to the other samples. For all samples, cells exceeding 20% of mitochondrial gene counts were also filtered out. We used DoubletFinder^70^ to identify and remove cells based on transcript content. The number of cells removed at each filtering step per sample is included in **Table S1**.

##### Normalization, integration, and clustering

Using Seurat, per-sample cell counts were merged per developmental stage and normalized using its default log-normalization strategy. The FindVariableFeatures parameter was set at 4,000. Integration was performed using the Seurat pipeline (v5.0.3),^69^ selecting anchors and using them to integrate the datasets. The k.weight parameter was set as default, with the exception of the neonatal germ cells due to low number of cells in the subset. After integration, the data was scaled and principal component analysis (PCA) was performed using the default Seurat parameters. Subsequently, we calculated the intrinsic dimensions of the PCA results using the maxLikGlobalDimEst() function of the IntrinsicDimension package^72^ (NM: 8; PPM: 14; PM: 10; AM: 12). The estimated intrinsic dimensions were rounded up or down and used as the dimension parameters for the RunUMAP() and FindNeighbors() commands respecting the default values for the remaining parameters (knn=20). Finally, we chose the resolution of the clustering using ClusTree,^73^ which analyzes the stability of the clustering at different resolutions. For the neonatal, pre-pubertal, pubertal, and adult datasets, we selected the lowest resolution with the highest stability (NM: 0.1; PPM: 0.1; PM: 0.1; AM: 0.1), since large transcriptional differences were expected among the testicular cell types.

##### Cluster identity in single-cell data

We labelled cell clusters with their likely cell identities based on the expression of previously reported marker genes.^6^ These labels comprised Leydig (*HSD17B3*, *STAR*), peritubular myoid (*DCN*, *ACTA2*, *CLEC3B*), Sertoli (*SOX9, AMH*), endothelial (*VWF*), perivascular (*NOTCH3*), immune (*CD45*), and germ cells (*DAZL*, *MAGEA4*, *SYCP3*). Distribution of cell counts across cell types and age groups is displayed in **Supplementary table 1**.

##### Differential gene expression analysis

Differential gene expression (DGE) analysis was performed using the Seurat functions FindAllMarkers(), for a general overview of the differentially expressed genes (DEGs) among all cells in a dataset, or FindMarkers(), to compare the differential expression directly between two (or more) clusters allowing the sequential approach. The logFC_threshold, the min.pct, and the min.diff.pct parameters were set at 1, 30%, and 20%, respectively. The test used was from the package MAST,^74^ which is tailored to identify differentially expressed genes from scRNA-seq experiments.

##### Gene set enrichment analysis

We computed overlaps with the GSEA molecular signatures database^75,76^ by uploading the top 500 DEGs onto the online tool to enrich for gene ontology terms associated with biological processes using a false recovery rate (FDR) q-value lower than 0.05. The top 20 or top 10 obtained gene ontology terms were included in this study for the general testicular datasets and the germ cell subset clusters, respectively.

##### Germ cell subsets

Neonatal, pre-pubertal, pubertal, and adult germ cells were subset and further analysed separately. We integrated the data and followed the same pipeline as for the general datasets. This approach identified further cells with lower RNA content, higher mitochondrial gene content, and combined expression of marker genes for different cell types. These cells were considered misclassified, low-quality cells and excluded from further analysis.

Further bioinformatic analyses were performed in the germ cell subsets of neonatal (599 cells), pre-pubertal (7,078 cells), pubertal (1,835 cells), and adult (15,982 cells) marmoset testis datasets. A spermatogonial subset was parsed out of the adult germ cells and subjected to the same analyses. The estimated intrinsic dimensions for these subsets were 14 for neonatal, 16 for pre-pubertal, 8 for pubertal, 13 for adult germ cells, and 12 for adult spermatogonia. To study the germ cells subsets, we investigated higher resolutions, keeping a high stability of clustering, to dissect transcriptional substates within germ cell types (neonatal: 0.8; pre-pubertal: 0.5; pubertal: 0.5; adult: 0.3; adult spermatogonia: 0.8).

##### Cell cycle analysis

Cell cycle phases were assigned following the Seurat protocol (v5.0.3).^69^ Briefly, a cell-cycle score was calculated based on the expression of S- or G2/M-specific genes, and expression of neither of them leads to G1 assignment.

##### Gene regulatory network analysis

Single-Cell rEgulatory Network Inference and Clustering (SCENIC)^24^ analysis was performed in the germ cell subsets to infer active gene regulatory networks. As input, we used the scRNA-seq expression matrices. We searched transcription factor binding motifs within 500 bp upstream or 10 kb centred around the transcription start site. The network activity was scored in each cell using AUCell^24^ and subsequently binarized into active or inactive regulons. The generated activity matrices were embedded into the germ cell Seurat objects to identify differentially active regulons specific to each germ cell cluster identity.

##### Pearson’s correlations

To perform Pearsońs correlations analysis between the germ cell subsets during development, we used the function AggregateExpression() from the Seurat package (v5.0.3)^69^ to generate RNA summed counts (“pseudobulk”) of each germ cell cluster. Using this RNA summed counts, we compared the transcriptomes of the germ cell clusters across developmental stages, calculating Pearsońs correlations coefficients between them. Peritubular myoid cells were included for comparison and showed the lowest correlation to the germ cell clusters both intra-subset and inter-subset, demonstrating that the similarities are not influenced by batch effect.

### QUANTIFICATION AND STATISTICAL ANALYSIS

To identify significant differences in the number of positive cells for MAGEA4, UTF1, PIWIL4, DPPA4, and NANOS3 per round tubule between neonatal, pre-pubertal, pubertal, and adult samples, normality was first assessed using the Shapiro-Wilk test. Normal distribution could not be discarded in any of the datasets (**Table S6**). Homoscedasticity of whole datasets was tested using the Levene’s test, and in every case, the results showed homogeneity in the variances. We performed ANOVA to compare between the four groups, with Tukey’s test as post-hoc test. For the morphological correlation to the marker-based classification of spermatogonial subpopulations, the same approach was conducted. The Shapiro-Wilk test was lower than 0.05 in the percentages of UTF1^+^ A_dark_ spermatogonia in pre-pubertal samples and in DPPA4^+^ A_dark_ spermatogonia in pre-pubertal and adult samples. For comparisons involving these data, we used the Kruskal-Wallis’ test and Dunn’s test with Bonferroni correction as post-hoc test.

All the statistical analyses were performed in R (version 4.4.1) and are shown in the Supplementary tables.

## ADDITIONAL RESOURCES

