## Supplementary material for "Spermatogonial stem cells and their differentiation roadmap throughout marmoset development": Key Resource Table

**Key resources table**

| REAGENT or RESOURCE | SOURCE | IDENTIFIER |
| --- | --- | --- |
| Antibodies | | |
| PE anti-marmoset CD45 Antibody (1:50) | BioLegend | Cat# 250204, RRID: AB_2562120 |
| Goat anti-human PGLYRP1 Antibody (1:100) | R&D System | Cat# AF2590, RRID: AB_442160 |
| Anti-CD163 antibody, Rabbit monoclonal (1:50) | Sigma-Aldrich | Cat# SAB5600260 |
| Anti-Ki67 antibody (1:50) | Abcam | Cat# ab15580, RRID: AB_443209 |
| Ki-67 (8D5) Mouse Monoclonal Antibody (1:50) | Cell Signaling | Cat# 9449, RRID: AB_2797703 |
| Mouse monoclonal anti-AP2γ (1:50) | Santa Cruz | Cat# sc12762, RRID: AB_667770 |
| Rabbit polyclonal anti-DAZL (1:100) | Abcam | Cat# ab34139, RRID: AB_731849 |
| Mouse monoclonal anti-OCT4 (1:50) | Santa Cruz | Cat# sc5279, RRID: AB_628051 |
| Rabbit polyclonal anti-LIN28A (1:100) | Cell Signaling | Cat# 3978, RRID: AB_2297060 |
| Mouse monoclonal anti-MAGEA4 (1:20) | Prof. Spagnoli; University of Basel, Switzerland | N/A |
| Rabbit polyclonal anti-PIWIL4 (1:100) | LSBio | Cat# LS-C482396, RRID: AB_2782961 |
| Anti-CITED2 antibody (1:100) | Abcam | Cat# ab184145 |
| Anti-DPPA4 Antibody (1:100) | Human Protein Atlas | Cat# HPA035250, RRID: AB_2674536 |
| Rabbit polyclonal anti-NANOS3 (1:50) | Sigma-Aldrich | Cat# HPA062989, RRID: AB_2684916 |
| Rabbit polyclonal anti-EGR1 (1:100) | Atlas Antibodies | Cat# HPA029937, RRID: AB_2673258 |
| RHOXF2B Polyclonal Antibody (1:500) | ThermoFisher | Cat# PA5-35225, RRID: AB_2552535 |
| Anti-UTF-1 Antibody, clone 5G10.2 (1:20) | Millipore | Cat# MAB4337, RRID: AB_827541 |
| IgG from mouse serum | Sigma-Aldrich | Cat# I5381, RRID: AB_1163670 |
| IgG from rabbit serum | Sigma-Aldrich | Cat# I5006, RRID: AB_1163659 |
| Goat F(ab’)2 Anti-Mouse IgG  - (Fab’)2 (Biotin) | Abcam | Cat# ab5886, RRID: AB_954791 |
| Goat F(ab’)2 Anti-Rabbit IgG  - H&L (Biotin) | Abcam | Cat# ab6012, RRID: AB_954766 |
| Alexa Fluor® 488 AffiniPure® F(ab')₂ Fragment Donkey Anti-Rabbit IgG (H+L) | Jackson  ImmunoResearch  Labs | Cat# 711-546-152, RRID: AB_2340619 |
| Donkey anti-Mouse IgG (H+L) Highly Cross-Adsorbed Secondary Antibody, Alexa Fluor™ 488 | ThermoFisher | Cat# A-21202, RRID: AB_141607 |
| Donkey anti-Mouse IgG (H+L) Highly Cross-Adsorbed Secondary Antibody, Alexa Fluor™ 647 | ThermoFisher | Cat# A-31571, RRID: AB_162542 |
| Donkey anti-Rabbit IgG (H+L) Highly Cross-Adsorbed Secondary Antibody, Alexa Fluor™ 488 | ThermoFisher | Cat# A-21206, RRID: AB_2535792 |
| Donkey anti-Rabbit IgG (H+L) Highly Cross-Adsorbed Secondary Antibody, Alexa Fluor™ 647 | ThermoFisher | Cat# A-31573, RRID: AB_2536183 |
| Donkey anti-Goat IgG (H+L) Cross-Adsorbed Secondary Antibody, Alexa Fluor™ 647 | ThermoFisher | Cat# A-21447, RRID: AB_2535864 |
| Biological samples | | |
| Fresh marmoset testicular tissue | Central Animal Facility of the University clinic Münster |  |
| Bouin’s-fixed paraffin-embedded marmoset testicular tissue | Tissue bank in the Center of Reproductive Medicine and Andrology |  |
| PFA-fixed paraffin-embedded marmoset testicular tissue | Tissue bank in the Center of Reproductive Medicine and Andrology |  |
| Chemicals, peptides, and recombinant proteins | | |
| Streptavidin - Peroxidase  from Streptomyces avidinii | Sigma-Aldrich | Cat# S5512 |
| Bovine serum albumin | Sigma-Aldrich | Cat# A9647 |
| Goat serum | Sigma-Aldrich | Cat# G6767 |
| 3,30-diaminobenzidine tetrahydrochloride | Applichem | Cat# A0596.0001 |
| Mayer´s hematoxylin | Sigma-Aldrich | Cat# MHS32-1L |
| Merckoglass liquid cover glass for microscopy | Sigma-Aldrich | Cat# 1039730001 |
| Glycine | Sigma-Aldrich | Cat# G7126 |
| Triton X-100 | Sigma-Aldrich | Cat# 93443 |
| Tween 20 | Sigma-Aldrich | Cat# 655205 |
| Donkey serum | Sigma-Aldrich | Cat# S30 |
| VectaShield Mounting Medium with DAPI | Vector Laboratories | Cat# H-1200 |
| Minimum Essential Medium Alpha | Gibco | Cat# 22561-021 |
| Collagenase | Sigma-Aldrich | Cat# C9891 |
| DNAse | Sigma-Aldrich | Cat# SLCJ6659 |
| Trypsin | Gibco | Cat# 25725-018 |
| Hank’s Balanced Salt Solution | Gibco | Cat# 14175-053 |
| Propidium Iodide | Sigma-Aldrich | Cat# R6513 |
| Critical commercial assays | | |
| FlexAble 2.0 CoraLite® Plus 488 Antibody Labeling Kit for Rabbit IgG1 | Proteintech | Cat# KFA501 |
| FlexAble 2.0 CoraLite® Plus 555 Antibody Labeling Kit for Rabbit IgG | Proteintech | Cat# KFA502 |
| FlexAble 2.0 CoraLite® Plus 647 Antibody Labeling Kit for Rabbit IgG | Proteintech | Cat# KFA503 |
| Chromium Single Cell Kit v2 chemistry | 10x Genomics | Cat# PN-1000009 |
| Chromium Single Cell Kit v3 chemistry | 10x Genomics | Cat# PN-1000127 |
| Deposited data | | |
| Single-cell RNA-sequencing datasets of neonatal, pre-pubertal, pubertal, and adult marmoset testicular tissues | This paper | GEO: GSE198470 |
| Software and algorithms | | |
| 10x Genomics Cell Ranger v3.1.0 and v6.0.2 | Zheng et al.^1^ | RRID: SCR_023221 |
| Seurat v5.1 | Hao et al.^2^ | RRID: SCR_016341 |
| DoubletFinder v2.0.4 | McGinnis et al.^3^ | RRID: SCR_018771 |
| SoupX v1.6.2 | Young et al.^4^ | RRID: SCR_019193 |
| IntrinsicDimension v1.2 | Johnsson et al.^5^ | N/A |
| ClusTree v0.5.1 | Zappia et al.^6^ | RRID: SCR_016293 |
| MAST v**1.36.0** | McDavid^7^ | RRID: SCR_016340 |
| SCENIC | Aibar et al.^8^ | RRID: SCR_017247 |
| Rstatix v0.7.2 | Kassambra^9^ | RRID: SCR_021240 |
| car v3.1-3 | Fox et al.^10^ | RRID: SCR_022137 |
| Dplyr v1.1.4 | Wickham et al.^11^ | RRID: SCR_016708 |
| ggplot2 v3.5.2 | Wickham^12^ | RRID: SCR_014601 |
| ViewPoint v | PreciPoint | N/A |
| QuPath v0.6.0 | Bankhead et al.^13^ | RRID: SCR_018257 |
| Openxlsx v4.2.8.1 | Schauberger and Walker^14^ | RRID: SCR_019185 |
| Other | | |
