## Supplemental figures for "Spermatogonial stem cells and their differentiation roadmap throughout marmoset development"

#### Figure S1

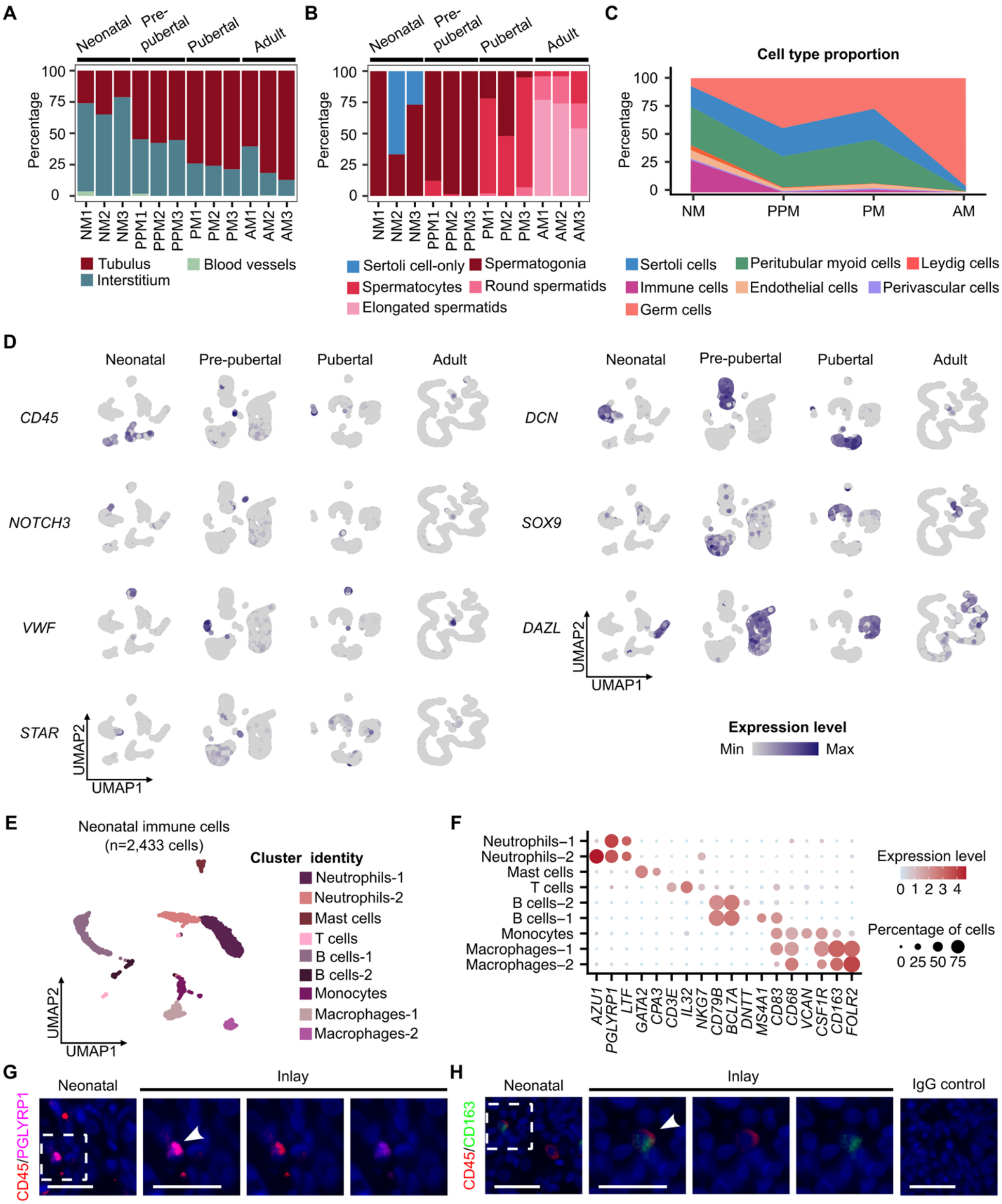
A
Neonatal
Pre-pubertal
Pubertal
Adult
Percentage
NM1 NM2 NM3 PPM1 PPM2 PPM3 PM1 PM2 PM3 AM1 AM2 AM3
Tubulus
Interstitium
Blood vessels
B
Sertoli cell-only
Spermatogonia
Spermatocytes
Round spermatids
Elongated spermatids
C
Cell type proportion
NM PPM PM AM
Sertoli cells
Peritubular myoid cells
Leydig cells
Immune cells
Endothelial cells
Perivascular cells
Germ cells
D
CD45
NOTCH3
VWF
STAR
DCN
SOX9
DAZL
UMAP1
UMAP2
Expression level
Min
Max
E
Neonatal immune cells (n=2,433 cells)
Cluster identity
Neutrophils-1
Neutrophils-2
Mast cells
T cells
B cells-1
B cells-2
Monocytes
Macrophages-1
Macrophages-2
F
AZU1 PGLYRP1 LTF GATA2 CPA3 CD3E IL32 NKG7 CD79B BCL7A DNTT MS4A1 CD83 CD68 VCAN CSF1R CD163 FOLR2
Expression level
Percentage of cells
0 25 50 75
G
CD45/PGLYRP1
Neonatal
Inlay
H
CD45/CD163
IgG control

#### Figure S2

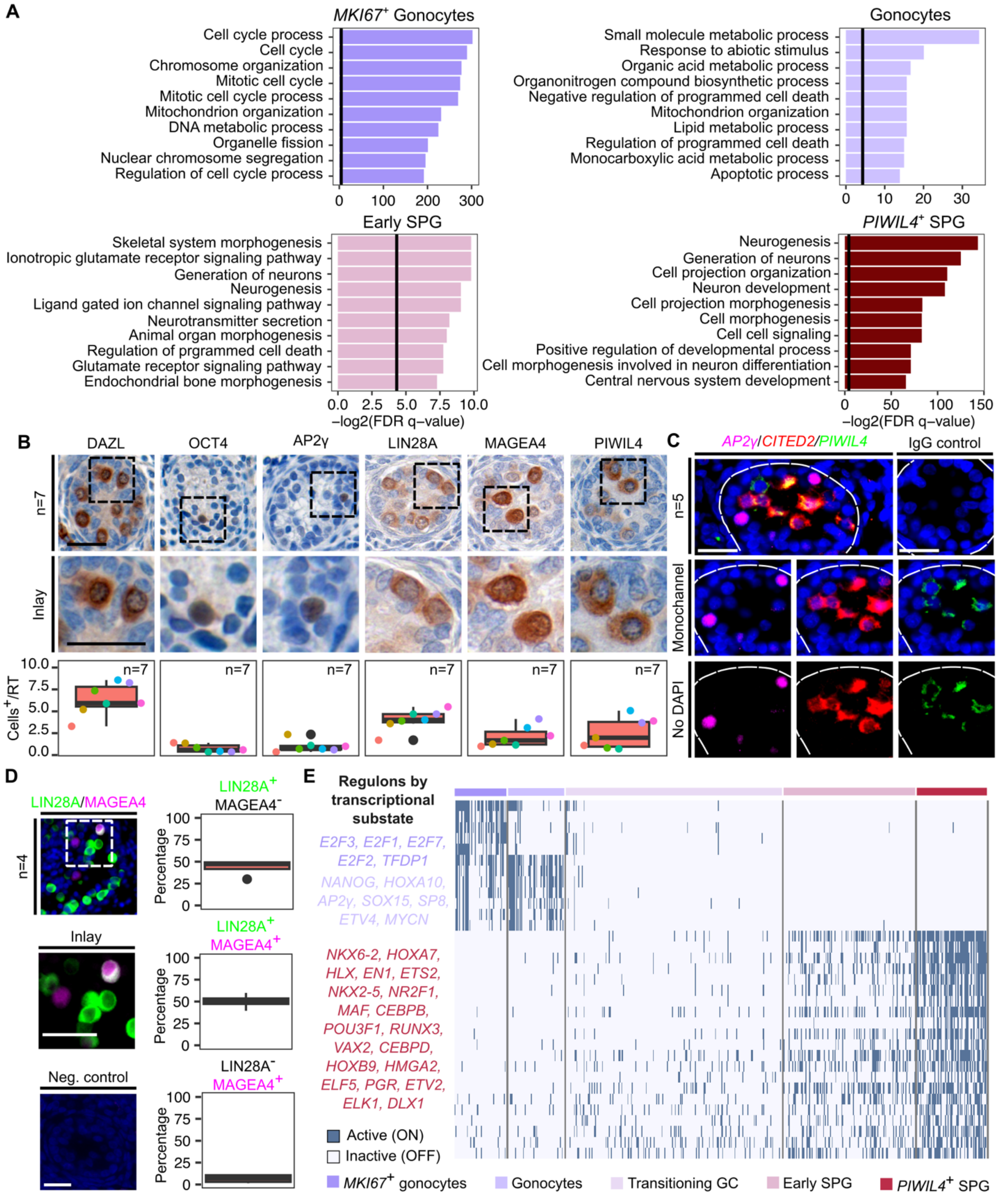
A
MKI67+ Gonocytes
Cell cycle process
Cell cycle
Chromosome organization
Mitotic cell cycle
Mitotic cell cycle process
Mitochondrion organization
DNA metabolic process
Organelle fission
Nuclear chromosome segregation
Regulation of cell cycle process
0 100 200 300
Gonocytes
Small molecule metabolic process
Response to abiotic stimulus
Organic acid metabolic process
Organonitrogen compound biosynthetic process
Negative regulation of programmed cell death
Mitochondrion organization
Lipid metabolic process
Regulation of programmed cell death
Monocarboxylic acid metabolic process
Apoptotic process
0 10 20 30
Early SPG
Skeletal system morphogenesis
Ionotropic glutamate receptor signaling pathway
Generation of neurons
Neurogenesis
Ligand gated ion channel signaling pathway
Neurotransmitter secretion
Animal organ morphogenesis
Regulation of prgrammed cell death
Glutamate receptor signaling pathway
Endochondrial bone morphogenesis
0.0 2.5 5.0 7.5 10.0
−log2(FDR q−value)
PIWIL4+ SPG
Neurogenesis
Generation of neurons
Cell projection organization
Neuron development
Cell projection morphogenesis
Cell morphogenesis
Cell cell signaling
Positive regulation of developmental process
Cell morphogenesis involved in neuron differentiation
Central nervous system development
0 50 100 150
−log2(FDR q−value)
B
DAZL OCT4 AP2γ LIN28A MAGEA4 PIWIL4
n=7
Inlay
Cells+/RT
10.0 7.5 5.0 2.5 0.0
n=7
C
AP2γ/CITED2/PIWIL4
IgG control
n=5
Monochannel
No DAPI
D
LIN28A/MAGEA4
n=4
Inlay
Neg. control
LIN28A+ MAGEA4−
LIN28A+ MAGEA4+
LIN28A− MAGEA4+
Percentage
100 75 50 25 0
E
Regulons by transcriptional substate
E2F3, E2F1, E2F7, E2F2, TFDP1
NANOG, HOXA10, AP2γ, SOX15, SP8, ETV4, MYCN
NKX6-2, HOXA7, HLX, EN1, ETS2, NKX2-5, NR2F1, MAF, CEBPB, POU3F1, RUNX3, VAX2, CEBPD, HOXB9, HMGA2, ELF5, PGR, ETV2, ELK1, DLX1
Active (ON)
Inactive (OFF)
MKI67+ gonocytes
Gonocytes
Transitioning GC
Early SPG
PIWIL4+ SPG

### Figure S3

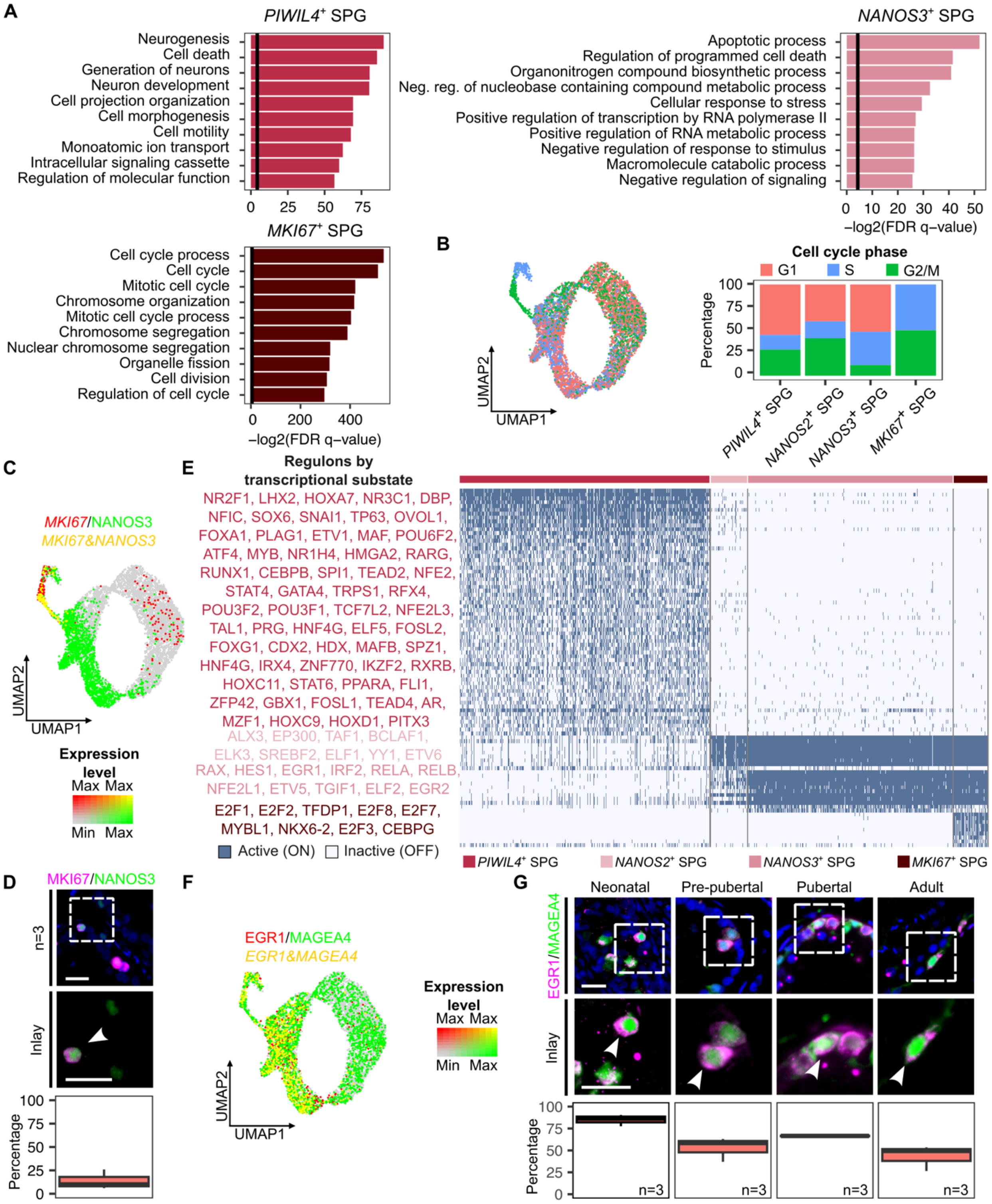

A
PIWIL4+ SPG
Neurogenesis
Cell death
Generation of neurons
Neuron development
Cell projection organization
Cell morphogenesis
Cell motility
Monoatomic ion transport
Intracellular signaling cassette
Regulation of molecular function
0 25 50 75
NANOS3+ SPG
Apoptotic process
Regulation of programmed cell death
Organonitrogen compound biosynthetic process
Neg. reg. of nucleobase containing compound metabolic process
Cellular response to stress
Positive regulation of transcription by RNA polymerase II
Positive regulation of RNA metabolic process
Negative regulation of response to stimulus
Macromolecule catabolic process
Negative regulation of signaling
0 10 20 30 40 50
−log2(FDR q−value)
MKI67+ SPG
Cell cycle process
Cell cycle
Mitotic cell cycle
Chromosome organization
Mitotic cell cycle process
Chromosome segregation
Nuclear chromosome segregation
Organelle fission
Cell division
Regulation of cell cycle
0 200 400
−log2(FDR q−value)
B
UMAP2
UMAP1
Cell cycle phase
G1 S G2/M
Percentage
100 75 50 25 0
PIWIL4+ SPG
NANOS2+ SPG
NANOS3+ SPG
MKI67+ SPG
C
MKI67/NANOS3
MKI67&NANOS3
UMAP2
UMAP1
Expression level
Max Max
Min Max
E
Regulons by transcriptional substate
NR2F1, LHX2, HOXA7, NR3C1, DBP, NFIC, SOX6, SNAI1, TP63, OVOL1, FOXA1, PLAG1, ETV1, MAF, POU6F2, ATF4, MYB, NR1H4, HMGA2, RARG, RUNX1, CEBPB, SPI1, TEAD2, NFE2, STAT4, GATA4, TRPS1, RFX4, POU3F2, POU3F1, TCF7L2, NFE2L3, TAL1, PRG, HNF4G, ELF5, FOSL2, FOXG1, CDX2, HDX, MAFB, SPZ1, HNF4G, IRX4, ZNF770, IKZF2, RXRB, HOXC11, STAT6, PPARA, FLI1, ZFP42, GBX1, FOSL1, TEAD4, AR, MZF1, HOXC9, HOXD1, PITX3
ALX3, EP300, TAF1, BCLAF1, ELK3, SREBF2, ELF1, YY1, ETV6
RAX, HES1, EGR1, IRF2, RELA, RELB, NFE2L1, ETV5, TGIF1, ELF2, EGR2
E2F1, E2F2, TFDP1, E2F8, E2F7, MYBL1, NKX6-2, E2F3, CEBPG
Active (ON) Inactive (OFF)
PIWIL4+ SPG
NANOS2+ SPG
NANOS3+ SPG
MKI67+ SPG
D
MKI67/NANOS3
n=3
Inlay
Percentage
100 75 50 25 0
F
EGR1/MAGEA4
EGR1&MAGEA4
UMAP2
UMAP1
Expression level
Max Max
Min Max
G
Neonatal
Pre-pubertal
Pubertal
Adult
EGR1/MAGEA4
Inlay
Percentage
100 75 50 25 0
n=3 n=3 n=3 n=3

#### Figure S4

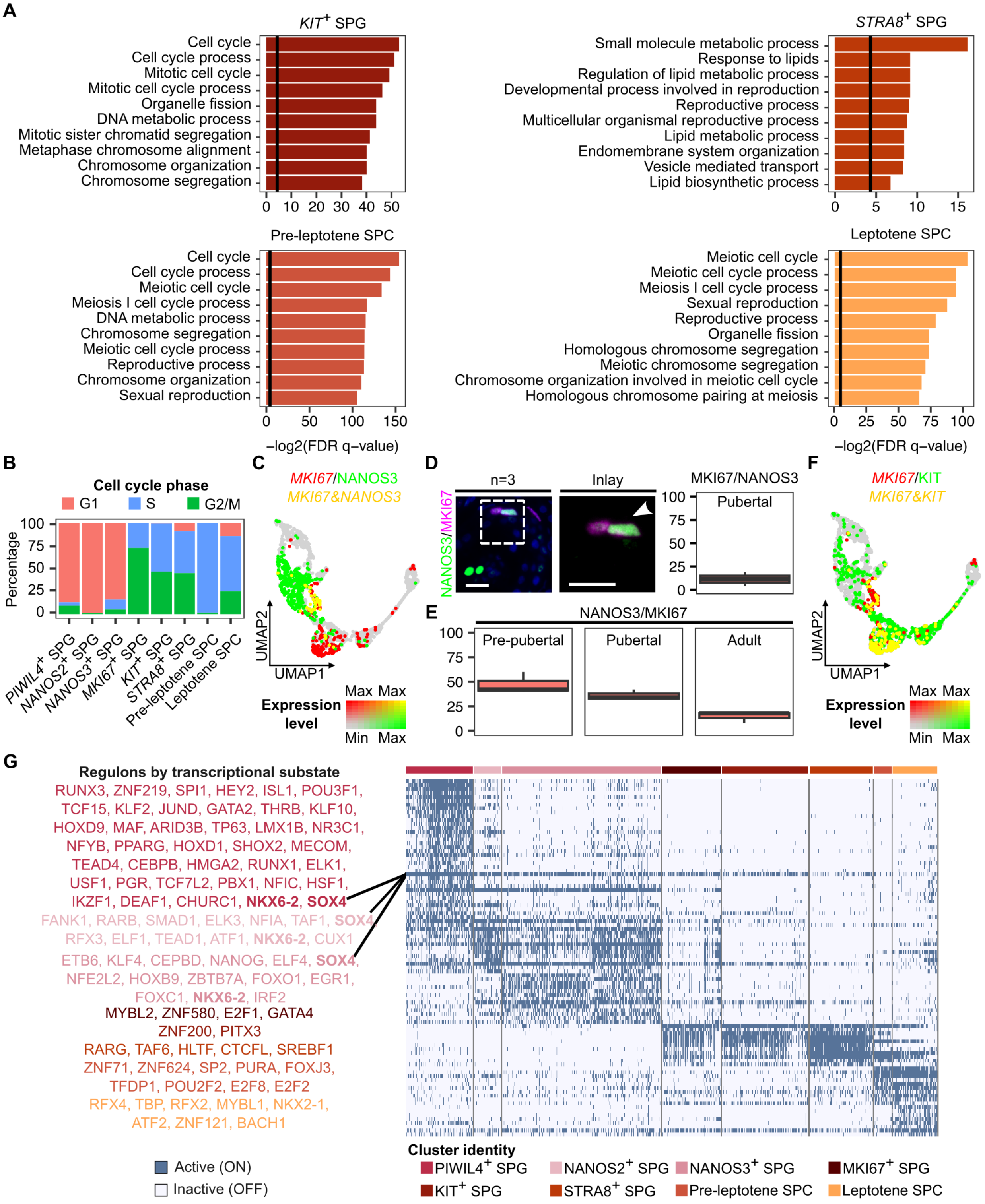
A
KIT+ SPG
Cell cycle
Cell cycle process
Mitotic cell cycle
Mitotic cell cycle process
Organelle fission
DNA metabolic process
Mitotic sister chromatid segregation
Metaphase chromosome alignment
Chromosome organization
Chromosome segregation
STRA8+ SPG
Small molecule metabolic process
Response to lipids
Regulation of lipid metabolic process
Developmental process involved in reproduction
Reproductive process
Multicellular organismal reproductive process
Lipid metabolic process
Endomembrane system organization
Vesicle mediated transport
Lipid biosynthetic process
Pre-leptotene SPC
Cell cycle
Cell cycle process
Meiotic cell cycle
Meiosis I cell cycle process
DNA metabolic process
Chromosome segregation
Meiotic cell cycle process
Reproductive process
Chromosome organization
Sexual reproduction
Leptotene SPC
Meiotic cell cycle
Meiotic cell cycle process
Meiosis I cell cycle process
Sexual reproduction
Reproductive process
Organelle fission
Homologous chromosome segregation
Meiotic chromosome segregation
Chromosome organization involved in meiotic cell cycle
Homologous chromosome pairing at meiosis
−log2(FDR q−value)
−log2(FDR q−value)
B
Cell cycle phase
G1
S
G2/M
Percentage
PIWIL4+ SPG
NANOS2+ SPG
NANOS3+ SPG
MKI67+ SPG
KIT+ SPG
STRA8+ SPG
Pre-leptotene SPC
Leptotene SPC
C
MKI67/NANOS3
MKI67&NANOS3
UMAP2
UMAP1
Expression level
Max Max
Min Max
D
n=3
Inlay
NANOS3/MKI67
MKI67/NANOS3
Pubertal
E
NANOS3/MKI67
Pre-pubertal
Pubertal
Adult
F
MKI67/KIT
MKI67&KIT
UMAP2
UMAP1
Expression level
Max Max
Min Max
G
Regulons by transcriptional substate
RUNX3, ZNF219, SPI1, HEY2, ISL1, POU3F1,
TCF15, KLF2, JUND, GATA2, THRB, KLF10,
HOXD9, MAF, ARID3B, TP63, LMX1B, NR3C1,
NFYB, PPARG, HOXD1, SHOX2, MECOM,
TEAD4, CEBPB, HMGA2, RUNX1, ELK1,
USF1, PGR, TCF7L2, PBX1, NFIC, HSF1,
IKZF1, DEAF1, CHURC1, NKX6-2, SOX4
FANK1, RARB, SMAD1, ELK3, NFIA, TAF1, SOX4
RFX3, ELF1, TEAD1, ATF1, NKX6-2, CUX1
ETB6, KLF4, CEPBD, NANOG, ELF4, SOX4,
NFE2L2, HOXB9, ZBTB7A, FOXO1, EGR1,
FOXC1, NKX6-2, IRF2
MYBL2, ZNF580, E2F1, GATA4
ZNF200, PITX3
RARG, TAF6, HLTF, CTCFL, SREBF1
ZNF71, ZNF624, SP2, PURA, FOXJ3,
TFDP1, POU2F2, E2F8, E2F2
RFX4, TBP, RFX2, MYBL1, NKX2-1,
ATF2, ZNF121, BACH1
Active (ON)
Inactive (OFF)
Cluster identity
PIWIL4+ SPG
NANOS2+ SPG
NANOS3+ SPG
MKI67+ SPG
KIT+ SPG
STRA8+ SPG
Pre-leptotene SPC
Leptotene SPC

### Figure S5

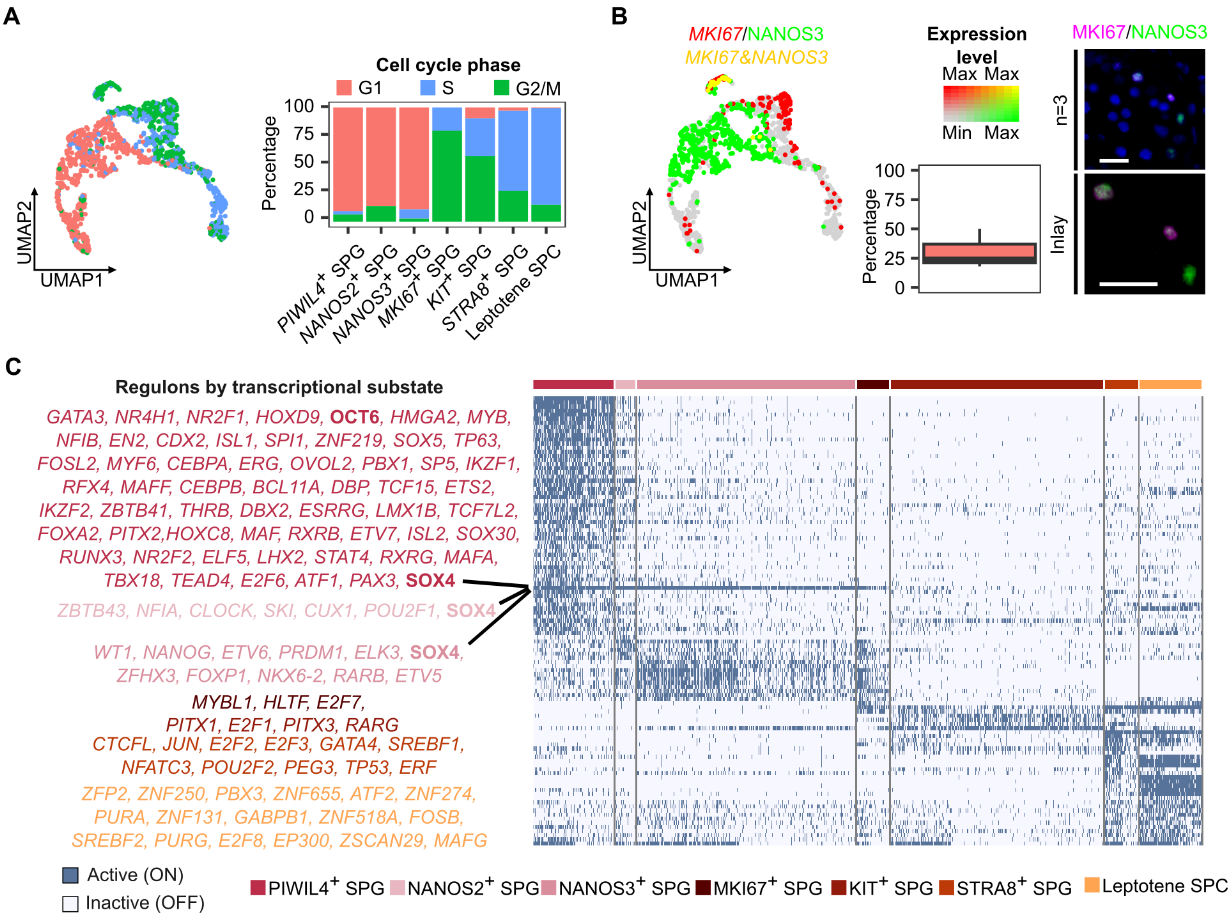
A
UMAP2
UMAP1
Cell cycle phase
G1
S
G2/M
Percentage
100
75
50
25
0
PIWIL4+ SPG
NANOS2+ SPG
NANOS3+ SPG
MKI67+ SPG
KIT+ SPG
STRA8+ SPG
Leptotene SPC
B
MKI67/NANOS3
MKI67&NANOS3
Expression level
Max Max
Min Max
UMAP2
UMAP1
Percentage
100
75
50
25
0
MKI67/NANOS3
n=3
Inlay
C
Regulons by transcriptional substate
GATA3, NR4H1, NR2F1, HOXD9, OCT6, HMGA2, MYB,
NFIB, EN2, CDX2, ISL1, SPI1, ZNF219, SOX5, TP63,
FOSL2, MYF6, CEBPA, ERG, OVOL2, PBX1, SP5, IKZF1,
RFX4, MAFF, CEBPB, BCL11A, DBP, TCF15, ETS2,
IKZF2, ZBTB41, THRB, DBX2, ESRRG, LMX1B, TCF7L2,
FOXA2, PITX2,HOXC8, MAF, RXRB, ETV7, ISL2, SOX30,
RUNX3, NR2F2, ELF5, LHX2, STAT4, RXRG, MAFA,
TBX18, TEAD4, E2F6, ATF1, PAX3, SOX4
ZBTB43, NFIA, CLOCK, SKI, CUX1, POU2F1, SOX4
WT1, NANOG, ETV6, PRDM1, ELK3, SOX4,
ZFHX3, FOXP1, NKX6-2, RARB, ETV5
MYBL1, HLTF, E2F7,
PITX1, E2F1, PITX3, RARG
CTCFL, JUN, E2F2, E2F3, GATA4, SREBF1,
NFATC3, POU2F2, PEG3, TP53, ERF
ZFP2, ZNF250, PBX3, ZNF655, ATF2, ZNF274,
PURA, ZNF131, GABPB1, ZNF518A, FOSB,
SREBF2, PURG, E2F8, EP300, ZSCAN29, MAFG
Active (ON)
Inactive (OFF)
PIWIL4+ SPG
NANOS2+ SPG
NANOS3+ SPG
MKI67+ SPG
KIT+ SPG
STRA8+ SPG
Leptotene SPC

**Figure S6**

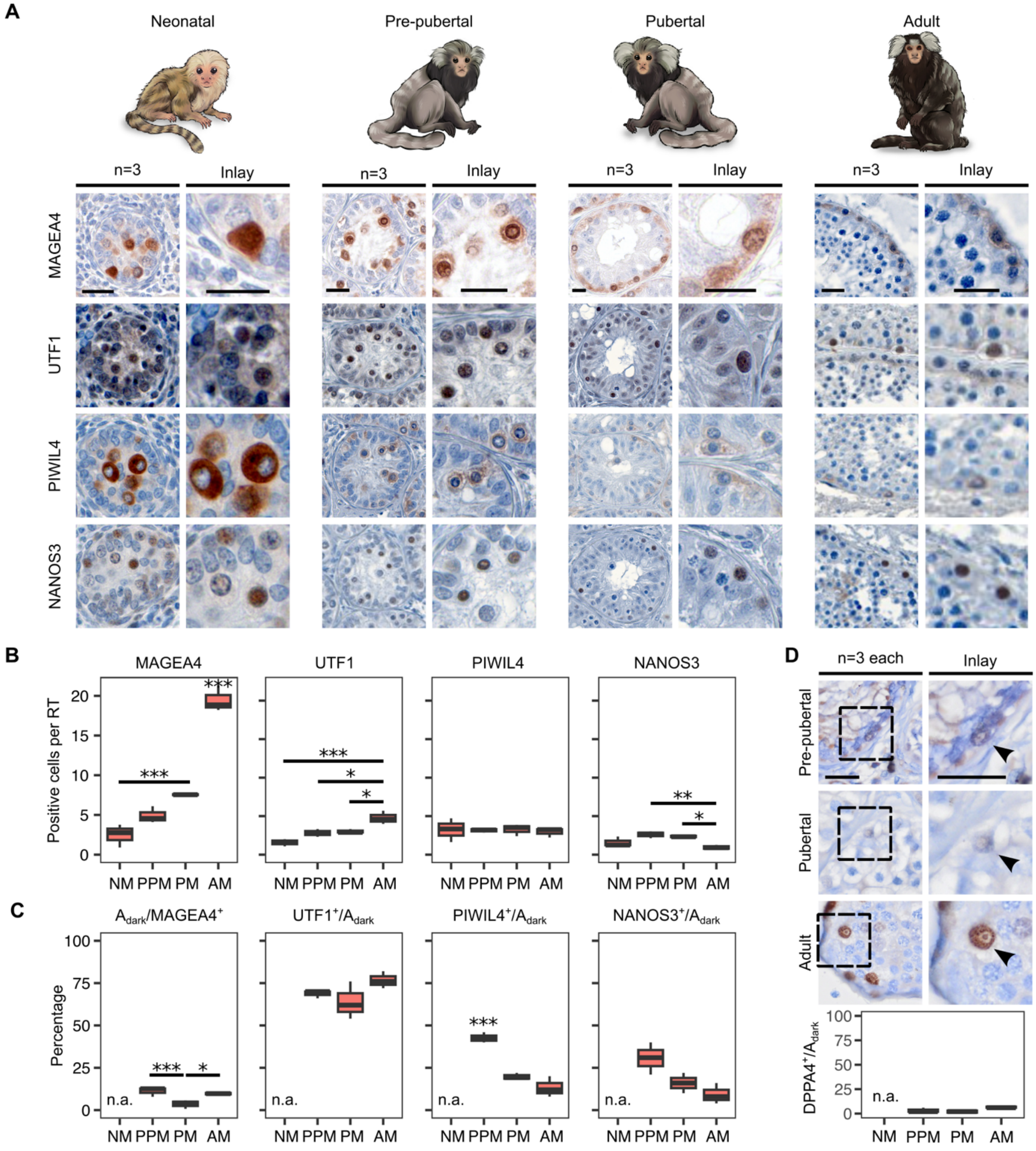
A
Neonatal
Pre-pubertal
Pubertal
Adult
n=3
Inlay
MAGEA4
UTF1
PIWIL4
NANOS3
B
MAGEA4
UTF1
PIWIL4
NANOS3
Positive cells per RT
NM PPM PM AM
C
Adark/MAGEA4+
UTF1+/Adark
PIWIL4+/Adark
NANOS3+/Adark
Percentage
n.a.
D
n=3 each
Inlay
Pre-pubertal
Pubertal
Adult
DPPA4+/Adark

#### Supplemental figure legends

Figure S1: **Tissue composition and somatic cell maturation (Related to Fig. 1)**

A) Barplots depicting the proportion of interstitium, blood vessels or seminiferous tubules determined by point counting per sample. Results are provided in **Table S1**.
B) Barplots showing the most advanced germ cell type per sample. Results are provided in **Table S1**.
C) Area plot portraying the changes in testicular cell type proportions per developmental stage. Results are provided in **Table S1**.
D) Feature plots showing expression of those marker genes used for cell cluster assignment.
E) Uniform manifold approximation and projection (UMAP) plot of the neonatal immune cell subset (n=2; 2,433 cells).
F) Dotplots showing the expression level of selected marker genes used for immune cell type assignment in the neonatal immune cell subset.
G) Representative immunofluorescence micrographs of neonatal marmoset testicular tissues stained for DAPI (blue), CD45 (red) and PGLYRP1 (magenta). Scale bars = 20 μm.
H) Representative immunofluorescence micrographs of neonatal marmoset testicular tissues stained for DAPI (blue), CD45 (red) and CD163 (green). Scale bars = 20 μm.

Figure S2: **Functional annotation and regulatory network activity in neonatal germ cells**

A) Gene ontology (GO) terms associated with the differentially expressed genes (DEGs) enriched in respective neonatal germ cell clusters, depicted as barplots. Bars depict the $–\log_2$ of the false discovery rate and the vertical line corresponds to FDRq-value = 0.05. No terms were obtained for Transitioning germ cells. Full lists of DEGs, GO terms and analyses are provided in **Table S2**.

B) Quantification results of the number of $DAZL^+$, $OCT4^+$, $AP2\gamma^+$, $LIN28A^+$, $MAGEA4^+$, and $PIWIL4^+$ cells per round tubule (RT). The upper row shows a representative micrograph of a round tubule for each marker, with their respective inlays in the row below. The lower row shows the quantification results neonatal marmoset testicular tissue samples (n=7), represented as boxplots. Each colored dot corresponds to the same individual per staining. Scale bars = 20 μm. Results are provided in **Table S2**.

C) Representative immunofluorescence micrographs of neonatal marmoset testicular tissues stained for AP2γ (magenta), CITED2 (red) and PIWIL4 (green), counterstained with DAPI (blue). The dashed circles delimit a seminiferous tubule. Scale bars = 20 μm.

D) Representative immunofluorescence micrographs (right column) of neonatal marmoset testicular tissues stained for LIN28A (green) and MAGEA4 (magenta), counterstained with DAPI (blue). The dashed square highlights the inlay area. Scale bars = 20 μm. The left column shows the percentage of $LIN28A^+$ (top row), $LIN28A^+MAGEA4^+$ (middle row) and $MAGEA4^+$ cells (bottom row), represented as boxplots. Results are provided in **Table S2**.

E) Heatmap showing the predicted differentially active regulons when comparing all neonatal germ cells clusters.

Figure S3: **Functional annotation and regulatory network activity in pre-pubertal germ cells**

A) GO terms associated with the DEGs enriched in respective pre-pubertal germ cell clusters, depicted as barplots. Bars depict the $-\log_2$ of the false discovery rate and the vertical line corresponds to FDRq-value = 0.05. No terms were obtained for *NANOS2*$^+$ spermatogonia. DEGs, full list of GO terms and analyses are provided in **Table S3**.

B) UMAP plot of pre-pubertal germ cells categorized based on cell cycle analysis into G1, S, or G2/M phases (left) and respective barplot showing the proportions per cluster (right). Results are provided in **Table S3**.

C) Double feature plot showing the expression of *MKI67* (red), *NANOS3* (green), or both (yellow) in the pre-pubertal germ cell subset.

D) Representative immunofluorescence micrographs of pre-pubertal marmoset testicular tissues stained for MKI67 (magenta) and NANOS3 (green) counterstained with DAPI are shown, with the dashed square highlighting the inlay area. Quantification results in pre-pubertal marmoset testicular tissues (n=3) are shown as percentage for MKI67$^+$/NANOS3$^+$ in a boxplot. Scale bars = 20 µm. Results are provided in **Table S3**.

E) Heatmap displaying the predicted differentially active regulons when comparing all pre-pubertal germ cells clusters.

F) Double feature plot showing the expression of *EGR1* (red), *MAGEA4* (green), or both (yellow) in the pre-pubertal germ cell subset.

G) Co-localization analysis of EGR1 (magenta) and MAGEA4 (green). The upper panel shows representative immunofluorescence micrographs of neonatal, pre-pubertal, pubertal and adult marmoset testicular tissue samples stained against EGR1 and MAGEA4, with their respective inlays in the middle row. White arrowheads indicate double positive cells. The dashed square highlights the inlay area. The quantification results shown as percentage of $EGR1^{+}$ in $MAGEA4^{+}$ cells is represented as a boxplot (bottom row). Scale bars = 20 μm. Results are provided in **Table S3**.

Figure S4: **Differentiation unleashed in the pubertal testis**

A) GO terms associated with the DEGs enriched between the pubertal germ cell clusters, depicted as barplots. Bars depict the $-\log_2$ of the false discovery rate and the vertical line corresponds to FDRq-value = 0.05. Full lists of DEGs, all GO terms and analyses are provided in **Table S4**.

B) Barplots depicting the proportion of cells in G1, S, or G2/M phase per cluster. Results are provided in **Table S4**.

C) Double feature plot showing the expression of *MKI67* (red), *NANOS3* (green), or both (yellow) in the pubertal germ cell subset.

D) Co-localization of NANOS3 (green) and MKI67 (magenta) in pubertal marmoset testicular tissue samples (n=2). The dashed square highlights the inlay area. The quantification results are displayed as percentage of $MKI67^+$ in $NANOS3^+$ cells in a boxplot. Scale bars = 20 μm. Results are provided in **Table S4**.

E) Quantification results shown as percentage of $NANOS3^+$ in $MKI67^+$ cells in pre-pubertal, pubertal and adult marmoset testicular tissues represented as boxplots (n=3 each). Results are provided in **Table S4**.

F) Double feature plot showing expression of *MKI67* (red), *KIT* (green), or both (yellow) in the pubertal germ cell subset.

H) Heatmap showing the predicted differentially active regulons when comparing all pubertal germ cells clusters.

Figure S5: **Full germ cell differentiation in the adult testis**

A) UMAP plot of the adult germ cells categorized based on cell cycle analysis results into G1, S, or G2/M phases (left) and respective barplot (right).

B) Double feature plot showing the expression of *MKI67* (proliferation marker gene; red), *NANOS3* (green), or both (proliferating *NANOS3*$^+$ spermatogonia; yellow) in the adult spermatogonial subset (left). On the right, co-localization analysis of MKI67 (magenta) and NANOS3 (green) is shown. The quantification results shown as percentage for $MKI67^+$/$NANOS3^+$ is represented as a boxplot. The staining was performed on adult marmoset testicular tissue samples (n=3). Scale bars = 20 μm. Results are provided in **Table S5**.

C) Heatmap showing the predicted differentially active regulons when comparing all adult spermatogonial clusters. Regulons are indicated on the left side and are ordered accordingly to the row orders.

Figure S6: **The molecular fingerprint of spermatogonial subpopulations throughout development**

A) Representative immunohistochemical micrographs of MAGEA4, UTF1, PIWIL4 and NANOS3 in neonatal, pre-pubertal, pubertal and adult testicular tissues (n=3 each). Scale bars = 20 μm and 10 μm in the inlays.

B) Quantification results of $MAGEA4^+$, $UTF1^+$, $PIWIL4^+$ or $NANOS3^+$ cells per RT at each developmental stage, represented as a boxplot. Results are provided in **Table S6**.

C) Quantification results of the percentage of $A_{dark}$ spermatogonia within $MAGEA4^+$ cells, and of $UTF1^+$, $PIWIL4^+$ or $NANOS3^+$ cells within the $A_{dark}$ spermatogonia at pre-pubertal, pubertal and adult stages, represented as boxplots. Results are provided in **Table S6**.

D) Representative immunohistochemical micrographs of DPPA4 stainings, highlighting $DPPA4^+$ $A_{dark}$ spermatogonia in pre-pubertal, pubertal and adult testicular tissues (n=3 each) and the respective quantification results of $DPPA4^+$ cells within the $A_{dark}$ spermatogonia at each developmental stage, represented as a boxplot. Scale bars = 20 μm and 10 μm in the inlays. Results are provided in **Table S6**.
